# Illuminating the Function of the Orphan Transporter, SLC22A10 in Humans and Other Primates

**DOI:** 10.1101/2023.08.08.552553

**Authors:** Sook Wah Yee, Luis Ferrández-Peral, Pol Alentorn, Claudia Fontsere, Merve Ceylan, Megan L. Koleske, Niklas Handin, Virginia M. Artegoitia, Giovanni Lara, Huan-Chieh Chien, Xujia Zhou, Jacques Dainat, Arthur Zalevsky, Andrej Sali, Colin M. Brand, John A. Capra, Per Artursson, John W. Newman, Tomas Marques-Bonet, Kathleen M. Giacomini

## Abstract

SLC22A10 is classified as an orphan transporter with unknown substrates and function. Here we describe the discovery of the substrate specificity and functional characteristics of SLC22A10. The human SLC22A10 tagged with green fluorescent protein was found to be absent from the plasma membrane, in contrast to the SLC22A10 orthologs found in great apes. Estradiol-17β-glucuronide accumulated in cells expressing great ape SLC22A10 orthologs (over 4-fold, p<0.001). In contrast, human SLC22A10 displayed no uptake function. Sequence alignments revealed two amino acid differences including a proline at position 220 of the human SLC22A10 and a leucine at the same position of great ape orthologs. Site-directed mutagenesis yielding the human SLC22A10-P220L produced a protein with excellent plasma membrane localization and associated uptake function. Neanderthal and Denisovan genomes show human-like sequences at proline 220 position, corroborating that SLC22A10 were rendered nonfunctional during hominin evolution after the divergence from the pan lineage (chimpanzees and bonobos). These findings demonstrate that human SLC22A10 is a unitary pseudogene and was inactivated by a missense mutation that is fixed in humans, whereas orthologs in great apes transport sex steroid conjugates.

## Introduction

About 30% of the members of the large Solute Carrier Superfamily in the human genome have no known substrate^1^, representing a major gap in understanding human biology. Deorphaning is the process of determining the function of a protein that has not yet been characterized. For deorphaning proteins in the SLC superfamily, which includes multi-membrane spanning transporters, phylogenetic analysis represents the first step for identifying the substrates of an orphan transporter. Other methods include metabolomic methods in cells or in knockout mice^2–4^. For example, the substrate of mouse SLC16A6, a transporter in the MonoCarboxylate Transporter Family, MCT7, was discovered through the analysis of amino acid transport in cell lines that overexpressed MCT7^5^.

There has been increasing interest in deorphaning solute carrier transporters due to the significant potential role of SLCs in human physiology^1, 6^. While some orphan genes in the SLC superfamily encode proteins which have evolved other functions and do not participate in transmembrane solute flux, it is more probable that these multi-membrane spanning proteins primarily serve as transporters. Therefore, efforts to deorphan SLC family members should include attempts to identify their endogenous substrates, ligands of the transporter that are translocated across biological membranes. In certain cases, the roles of transporters have been discovered, yet the substrate of the transporter remains unknown. This creates gaps in our mechanistic understanding of the transporter’s function in those processes. For example, Ren *et al.* used untargeted metabolomics and found elevated levels of lipid diacylglycerol and altered fatty acid metabolites in liver and plasma samples of Mct6 knockout mice^7^. This finding supports a role of SLC16A5 in lipid and amino acid homeostasis, but does not reveal its substrates and as such, the mechanism remains poorly understood. Similarly, the function of the orphan transporter SLC38A10 (SNAT10) was assessed by studying mice lacking the *Slc38a10* gene (Slc38a10-deficient mice). The findings indicated that *Slc38a10*-deficient mice exhibited reduced body weight and lower plasma levels of threonine and histidine. However, no studies have specifically investigated whether these amino acids serve as substrates for SLC38A10^8^; therefore a gap in understanding the mechanism by which the transporter affects body weight remains. Information regarding recently deorphaned transporters is presented in a recent review^9^.

Orphan transporters can be found in over 20 families in the SLC superfamily^1^. In the Solute Carrier 22 family A (SLC22A), there are 28 members that transport organic ions and another 10 that are orphans^6^. Largely representing plasma membrane transporters, members of the SLC22A family are clustered together based on their charge specificity for organic cations (OCTs), organic anions (OATs), and organic zwitterion/cations (OCTNs). Solute carrier 22 family member 10 (SLC22A10) and its direct species orthologs are orphan transporters whose substrates and transport mechanisms are yet to be characterized. In humans, SLC22A10 has been given a protein name of OAT5. Based on Northern blotting^10^ and RNA seq studies (https://www.proteinatlas.org/ENSG00000184999-SLC22A10/tissue)^11^, human SLC22A10 is expressed specifically in the liver.

Orthologs of human SLC22A10 are present in some primates including great apes (https://useast.ensembl.org/Homo_sapiens/Gene/Compara_Ortholog?db=core;g=ENSG00000184999;r=11:63268022-63311783). Intrigued by this observation, our study aimed to identify the substrates of SLC22A10 and the transport mechanism by expressing primate orthologs of SLC22A10 in cell lines and performing analytical procedures including cellular uptake studies, metabolomic analyses and proteomic assays. We attempted to identify crucial amino acids that contribute to the differences in function between direct species orthologs in humans and great apes. Kinetic parameters and transport mechanisms of various predicted isoforms of SLC22A10 were determined, along with their ability to accumulate different endogenous ligands. Proteomic studies in cell lines recombinantly expressing human and chimpanzee SLC22A10 were conducted. To the best of our knowledge, this is the first study to characterize the function of the orphan transporter SLC22A10. Our study shows that human SLC22A10 was inactivated by a single missense mutation and is a unitary pseudogene. The ORF-disrupting mutation in SLC22A10, which led to Pro220, is not observed in great apes and primates. This particular amino acid is crucial for protein abundance and expression on the plasma membrane. Our work provides a roadmap for how orthologous genes, along with sequence comparison, and proteomic and transporter assays, can be used to deorphan the function of solute carrier proteins. These discoveries have significant evolutionary implications.

## Results

### Human SLC22A10 showed no expression on the plasma membrane and no transporter activity of prototypical anionic substrates of SLC22 family members

Human SLC22A10 is in a cluster that includes known organic anion transporters: SLC22A24 is closest, followed by SLC22A9, SLC22A11 and SLC22A12 (***Fig. 1A***). Phylogenetic analyses reveals that its closest homolog is SLC22A24. The substrates of SLC22A24 are steroid conjugates, bile acids and dicarboxylic acids, which our laboratory has successfully deorphaned^3^. Overexpression of human SLC22A10 tagged with GFP in the N-terminal resulted in no detection of a GFP-tagged protein on the plasma membrane (***Fig. 1B***). Furthermore, no uptake of prototypical organic anions was observed in cells expressing SLC22A10 whereas significant uptake was observed in cells expressing known SLC22 organic anion transporters including SLC22A6, SLC22A8 and SLC22A24 (***Fig. 1C***).

**Figure 1.**
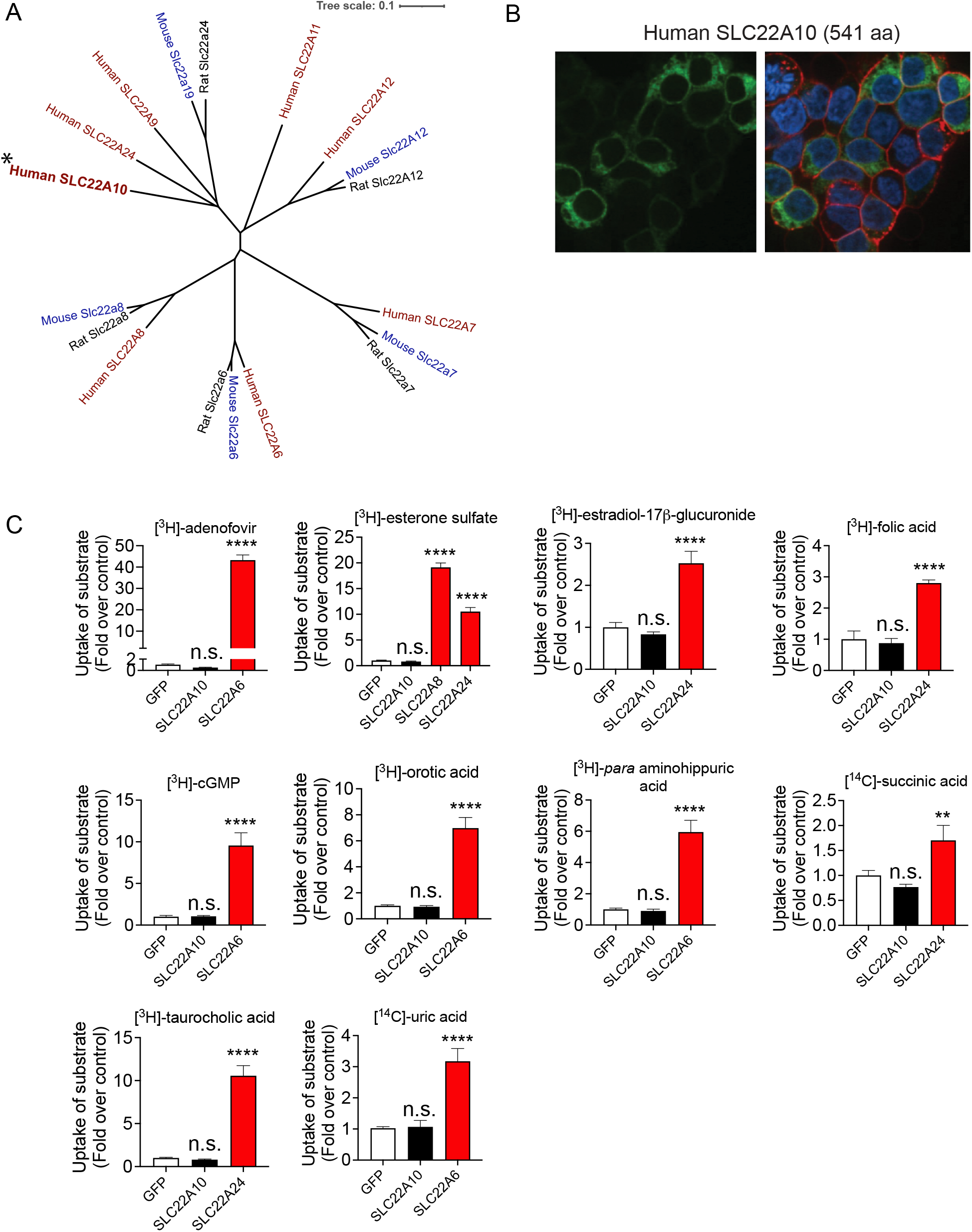
Analysis of the phylogenetic tree, plasma membrane expression of SLC22A10, and uptake of organic anion substrates of the human SLC22 family. **A.** Multiple sequence alignments were performed with reference amino acid sequences for each anion transporter from humans and rodents, using the Clustal Omega Multiple Sequence Alignment program (https://www.ebi.ac.uk/Tools/msa/clustalo/.) The dendrogram was generated from the output of the Clustal Omega alignment. **B.** Localization of human SLC22A10 conjugated to green fluorescent protein (GFP) was examined in HEK293 cells using high-content imaging and cellular staining with the plasma membrane marker wheat germ agglutinin (WGA). The results showed no colocalization of GFP-tagged SLC22A10 with WGA. **C.** Uptake of various radiolabeled organic anions, which are typical substrates of organic anion transporters in the SLC22A family, was assessed. Uptake was performed 48 hours after transient transfection of plasmids encoding human SLC22A10, GFP expression vector, and one other member in the SLC22A family as a positive control. Accumulation of substrates inside cells was determined after 15 minutes. Figure shows a representative plot from one experiment (mean ± S.D. from three replicate wells). The experiments were repeated at least one time and showed similar results. Multiple comparisons using one-way analysis of variance followed by Dunnett’s two-tailed test were performed. HEK293 cells transiently transfected with the GFP vector served as the control. The fold uptake of the substrate, relative to the control cells, was plotted based on one representative experiment conducted in triplicate wells (mean ± s.d.). The statistical significance for all the cells transfected with organic anion transporters SLC22A6, SLC22A8, or SLC2224 is p<0.001.

### The long isoform of chimpanzee and gorilla SLC22A10 was expressed on the plasma membrane whereas the short isoform was not

The organic anion transporters depicted in ***Fig. 1A*** consist of 536 to 563 amino acid proteins. Predictions from reliable sources such as Uniprot, Ensembl, and NCBI Nucleotide databases confirmed that the human SLC22A10 gene produces a 541 amino acid isoform. Conversely, according to reports from Ensembl and UniProt, the orthologs of SLC22A10 found in great apes are predicted to have two isoforms: a short isoform comprising 540 amino acids and a longer isoform containing 552 amino acids. Because there was no detectable expression of the human SLC22A10 on the plasma membrane of cells recombinantly expressing the transporter (***Fig. 1B***), we inquired whether the direct species orthologs in great apes exhibited a similar lack of plasma membrane expression when expressed recombinantly in cells. In fact, we observed that the long isoforms (552 amino acids) of both chimpanzee and gorilla SLC22A10 were detected on the plasma membrane (***Fig. 2A***). The shorter isoforms (540 amino acids) of chimpanzee, bonobo and gorilla SLC22A10 showed a similar lack of plasma membrane localization as the human ortholog, which consists of 541 amino acids (***Fig. 2A*** and ***Fig. 1B)*.**

**Figure 2.**
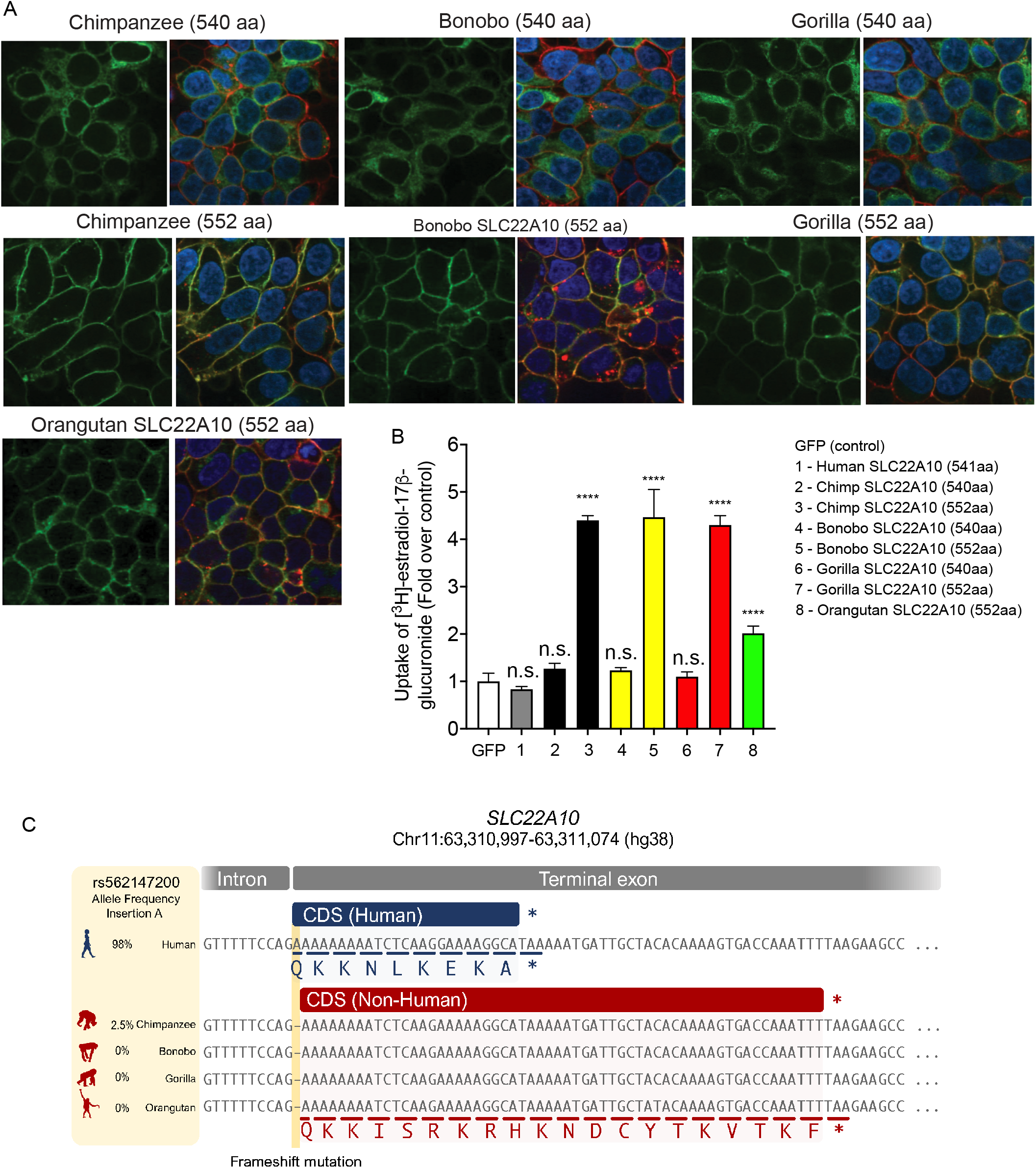
Localization to the plasma membrane, uptake, and sequence comparison of human SLC22A10 were examined in comparison with SLC22A10 from great apes (chimpanzee, bonobo, gorilla and orangutan). **A.** This figure shows the plasma membrane localization of SLC22A10 orthologs from great apes, which were conjugated to green fluorescent protein (GFP) in HEK293 cells. The GFP tag is located at the N-terminus of SLC22A10. Confocal imaging revealed that the 552 amino acid isoforms of SLC22A10 from chimpanzee, bonobo, gorilla, and orangutan primarily colocalized with wheat germ agglutinin (WGA) on the plasma membrane of the cell. In contrast, the 540 amino acid isoform of SLC22A10 from bonobo, chimpanzee, and gorilla showed no colocalization of GFP-tagged SLC22A10 with WGA on the plasma membrane, suggesting intracellular localization in the cytoplasm. **B.** The uptake of [^3^H]-estradiol-17b-glucuronide was determined in HEK293 cells overexpressing either a GFP expression vector or SLC22A10 expression vectors containing sequences from various primates including human, chimpanzee, bonobo, gorilla, and orangutan. SLC22A10 orthologs from chimpanzee, bonobo, gorilla, and orangutan expressing the longer isoform (552 amino acids) significantly accumulated [^3^H]-estradiol-17b-glucuronide. Please refer to the “Statistical Analysis” section for details on the statistical methods used to determine the significance of each cell transfected with the different SLC22A10 orthologs. **C.** Sequence alignments of the last exon of SLC22A10 in human, chimpanzee, bonobo, gorilla, and orangutan are shown. In humans, the frequency of the A-allele insertion is significantly greater (98%) than in chimpanzees (2.5%) and is not present in available sequences from bonobos, gorillas, or orangutans. The A-allele insertion results in the expression of human SLC22A10 with 541 amino acids, while bonobo, gorilla, orangutan and the majority of chimpanzees are predicted to express isoforms of SLC22A10 with 552 amino acids.

### Chimpanzee and gorilla SLC22A10 expressing the long isoform transport estradiol glucuronide but not other anions that are canonical substrates of members in the SLC22A family

Because the long forms of the great ape SLC22A10 showed a plasma membrane localization, we attempted to identify substrates of SLC22A10 using isotopic uptake assays in cells recombinantly expressing the long isoforms of the great ape transporters. Typical anions that are canonical substrates of members in the SLC22A family were screened for accumulation in human, chimpanzee, bonobo and gorilla expressing the long as well as the short isoforms. Significant accumulation of [^3^H]-estradiol-17β-glucuronide and [^3^H]-androstanediol-3α-glucuronide were observed in cells expressing chimpanzee and gorilla SLC22A10 encoding the long but not the short isoforms (***Fig. 2B***, ***Supplemental Fig. 1A***). No significant uptake was detected for other anions that are canonical substrates of members of the SLC22A family, such as estrone sulfate, taurocholic acid, cGMP, uric acid and succinic acid (***Supplemental Fig. 1A* to *F***). However, there was a small but significant uptake of [^3^H]-methotrexate in HEK293 cells expressing chimpanzee and gorilla SLC22A10 long isoforms (***Supplemental Fig. 1G***).

### The SLC22A10 protein in humans consists of 541 amino acids, resulting from a single nucleotide insertion that causes a frameshift in the last exon

The genetic mechanism that led to the formation of the 541 amino acid SLC22A10 protein in humans was investigated. Sequence alignments of the last exon (exon 10) of the SLC22A10 gene was compared between humans and great apes and revealed an insertion of one nucleotide leading to the expression of different isoforms in each species (see ***Fig. 2C***). In particular, humans exhibit an A nucleotide insertion at the first base pair of exon 10, which is highly prevalent with an allele frequency of 98% in all populations in gnomAD. In contrast, the adenosine insertion has a 2.5% allele frequency in chimpanzees and is not present in other great apes. The adenosine insertion in the human SLC22A10 gene causes a frameshift and results in a 541 amino acid protein, instead of the predicted 552 amino acids in the SLC22A10 gene of great apes and in the humans who do not harbor the adenosine insertion (***Fig. 2C***). We utilized our previously generated human and liver RNAseq data^12^ alongside a large human RNA-seq databases (recount3^13^) to validate the splicing event in human and chimpanzee SLC22A10. Our analysis confirmed that splicing mostly occurs at the exact orthologous genomic region in both species, utilizing the canonical splice sites in their corresponding genomes. Thus, we found no coordinated splicing alterations that compensate for the A nucleotide insertion. Consequently, the additional A nucleotide remains present in the final transcribed transcript in humans. This additional nucleotide provides evidence that human SLC22A10 protein contains 541 amino acids, while the chimpanzee SLC22A10 protein comprises 552 amino acids.

### Mutagenesis of a single amino acid, at position 220 of the human SLC22A10 ortholog, to the respective amino acid in great apes rescues the function of human SLC22A10

The alignment of human SLC22A10 and primate ortholog sequences revealed differences in amino acid positions p.Met18IIe and p.Pro220Leu (***Fig. 3A***). Interestingly, site-directed mutagenesis experiments demonstrated that the substitution of proline with leucine at position 220 (p.Pro220Leu) restored the plasma membrane localization and function of human SLC22A10, but the substitution of methionine with isoleucine at position 18 (p.Met18Ile) did not have any effect (see ***Fig. 3B***). In contrast, the replacement of leucine with proline at position 220 (p.Leu220Pro) abolished the localization and function of the chimpanzee SLC22A10 (see ***Fig. 3B*** and ***3C***). Additionally, we observed that a human-chimpanzee chimera protein, consisting of a fusion of human SLC22A10 (1-533) with chimpanzee SLC22A10 (534-552), while retaining the proline residue at position 220, showed no function (***Fig. 3B*** *and **3C***). However, with a leucine substitution at position 220, SLC22A10 remained functional (***Fig. 3B***).

**Figure 3.**
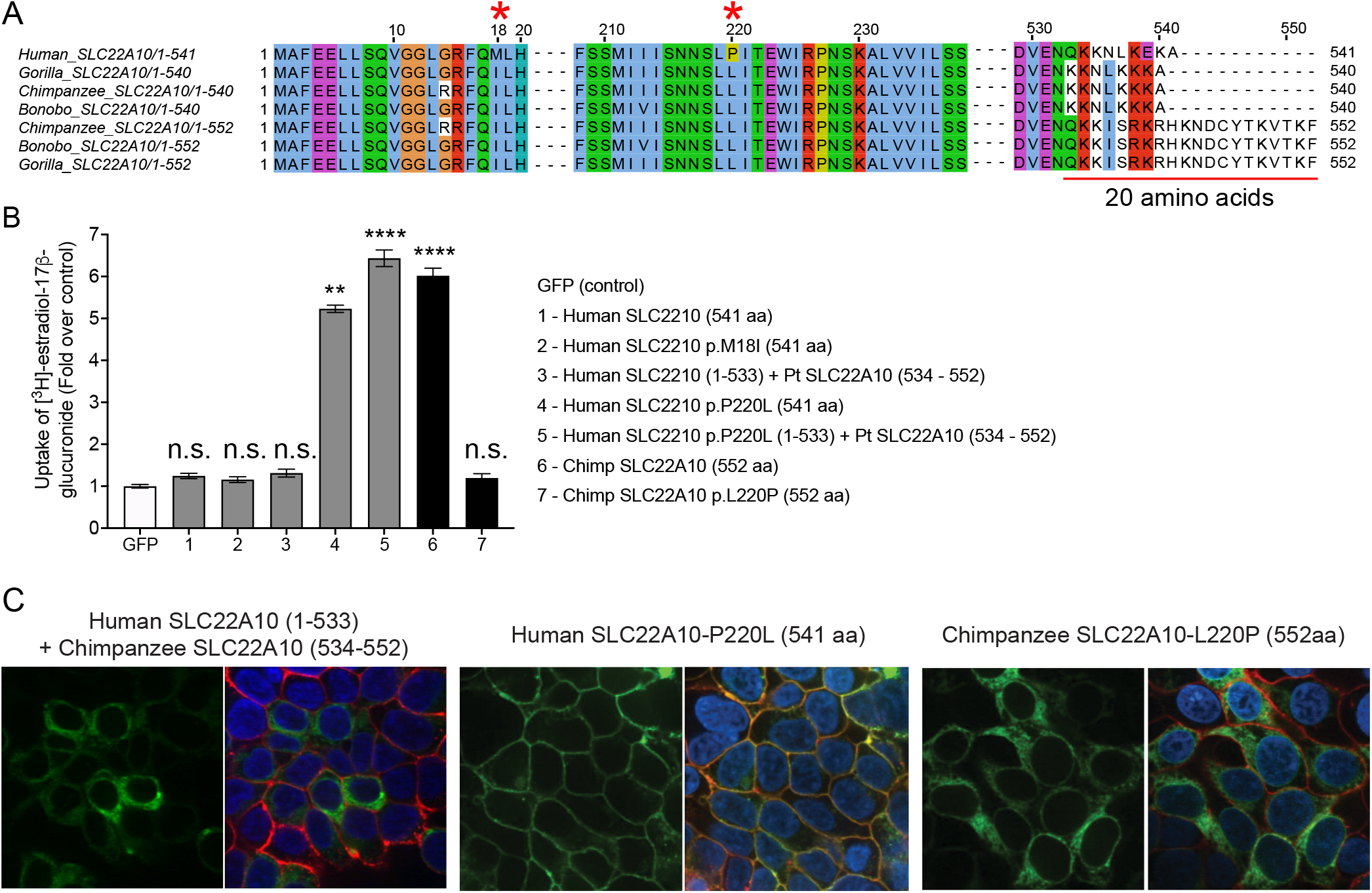
A single mutation of proline to leucine at amino acid position 220 of human SLC22A10 significantly enhances the accumulation of [^3^H]-estradiol-17b-glucuronide in HEK293. **A.** The amino acid sequence alignment of human SLC22A10 and SLC22A10 from other great apes (chimpanzee, bonobo and gorilla) shows that only the amino acids at positions 18 and 220 differ between the human ortholog and orthologs from great apes. Additionally, there are several amino acid differences starting at position 533. **B.** The uptake of [^3^H]-estradiol-17b-glucuronide in HEK293 cells transiently transfected with plasmids encoding human SLC22A10 with reference amino acids or amino acids that are similar to those found in other great apes, namely SLC22A10-p.M18I and SLC22A10-p.P220L. A chimeric protein consisting of the first 533 amino acids of human SLC22A10 and the last 19 amino acids of chimpanzee SLC22A10 (534-552) was also evaluated, but did not significantly accumulate [^3^H]-estradiol-17b-glucuronide compared to the chimeric protein with p.P220L. The fold uptake of the substrate, relative to the control (GFP) cells, was plotted based on one representative experiment conducted in triplicate wells (mean ± s.d.). The statistical significance for cells transfected with SLC22A10 #4 (Human SLC2210 p.P220L (541 aa)), #5 (Human SLC2210 p.P220L (1-533) + Pt SLC22A10 (534 - 552)) and #6 (Chimp SLC2210 (552 aa)) is p<0.001. **C.** This figure shows the plasma membrane localization of SLC22A10 conjugated to green fluorescent protein (GFP) in HEK293 cells. The GFP tag is located at the N-terminus of SLC22A10. Confocal imaging revealed that human SLC22A10-p.P220L localizes primarily to the plasma membrane of the cell, while there was no localization to the plasma membrane in cells expressing a chimeric protein or chimpanzee SLC22A10 with proline at the 220 amino acid position.

### Human and chimpanzee SLC22A10 with Pro220 exhibit lower protein expression compared to orthologs with Leu220

The objective of this study was to analyze the protein expression of SLC22A10 in HEK293 Flp-In cells that were transfected with either vector only, or the cDNA of human or chimpanzee SLC22A10. This was achieved by quantifying the global proteomes of the cells, with a specific focus on amino acids at position 220 of SLC22A10. Comparable transcript levels of SLC22A10 in HEK293 cells that over-expressed either human or chimpanzee SLC22A10, as well as the respective variants (p.P220L or p.L220P), were observed (**Supplemental Fig. 2**). However, as illustrated in **Table 1**, lower protein expression levels for the human SLC22A10 reference (Proline220) and chimpanzee SLC22A10-L220P were observed when compared to SLC22A10 with leucine at the 220 amino acid position. The results showed that protein levels of human SLC22A10 are approximately 10-fold lower in cells expressing human SLC22A10-Pro220 compared to human SLC22A10-Leu220 (**Table 1**) suggesting that human SLC22A10 is transcribed but the protein is unstable. The lower overall protein expression may explain the lack of detectable expression of human SLC22A10 on the plasma membrane in contrast to orthologs (both human and chimpanzee) of SLC22A10 with Leu220. In particular, human SLC22A10 was not detected on the plasma membrane (***Fig. 1B***), whereas the mutant, SLC22A10-Leu220 exhibited expression on the plasma membrane (***Fig. 3C***).

**Table 1.** The protein levels of SLC22A10 in HEK293 Flp-In cells that were transfected with expression vectors containing open reading frames of SLC22A10 orthologs. The protein levels are reported in fmol/µg protein and the values are the mean ± standard deviation from two independent experiments. The proteomic analysis was conducted using UniProt IDs to identify and characterize the detected SLC22A10 protein in the sample. Specifically, the SLC22A10 protein levels were searched against the UniProt database to find matches with the SLC22A10 protein sequence.

| **HEK293 Flp-In Cells** | **\*Human SLC22A10 protein (fmol/μg protein)** | **Other SLC22A10 isoforms (see UniprotID below)** |
| --- | --- | --- |
| Flp-In only | 0.00 | 0.01 ± 0.014 |
| Chimpanzee SLC22A10 (552 amino acid) | 0.00 | 10.23 ± 4.23 |
| Chimpanzee SLC22A10-L220P (552 amino acid) | 0.00 | 2.1 ± 1.64 |
| Human SLC22A10 (541 amino acid) | 0.083 ± 0.052 | 1.59 ± 1.51 |
| Human SLC22A10-P220L (541 amino acid) | 1.00 ± 0.21 | 14.58 ± 8.99 |

**\*Protein ID: Q63ZE4**
Uniprot Protein IDs for human and chimpanzee SLC22A10 protein sequence:
Human SLC22A10 (541 aa) - Q63ZE4
Chimpanzee SLC22A10 (540 aa) - A0A2I3TWY5
Chimpanzee SLC22A10 (552 aa) - A0A2J8LDY1
Human SLC22A10 (205 aa) - E9PIT2
Human SLC22A10 (264 aa) - E9PJB1
Human SLC22A10 (372 aa) - E9PMM0
Human SLC22A10 (155 aa) A0A804HHY1

### Human SLC22A10 with Pro220 is predicted to have poor stability compared to orthologs with Leu220

We tested the effect of the P220L mutation on the protein with the stability prediction pipeline tuned for transmembrane proteins^14^. The Rosetta physics-based score showed that P220L is a stabilizing mutation with a relatively low ΔΔG = -9.84 Rosetta Energy Units (REU), while the rank-normalized evolutionary-based ΔΔE scores of 0.0 and 0.16 for leucine and proline respectively also shows that leucine is more tolerated in this position than proline. The combination of both low ΔΔE and high ΔΔG is likely to cause loss of function via loss of stability and cellular abundance^15^.

### Analyses of SLC22A10 isoforms in chimpanzee, bonobo, orangutan and gibbon

Upon examining various databases (UniProt, Ensembl, and NCBI nucleotide) that predict SLC22A10 isoforms, we observed that chimpanzee, bonobo, orangutan, and gibbon are predicted to have different isoforms, including (i) chimpanzee (XM_024347215.1, 533 amino acid); (ii) bonobo (XM_034932815.1, 538 amino acid); (iii) gibbon (XM_003274123.1, 533 amino acid); and (iv) orangutan (XM_024255628.1, 533 amino acid). The shorter isoforms are derived from alternative acceptor sites (chimpanzee and bonobo) and exon extensions (orangutan and gibbon) resulting in 533-amino acid proteins (see ***Supplementary Fig. 3***). We conducted experiments using HEK293 Flp-In cells that were stably transfected with GFP-conjugated chimpanzee, bonobo, and gorilla SLC22A10 to examine the localization and function of the shorter isoforms. Our results indicated that the SLC22A10 533 amino acid isoform from chimpanzee, orangutan and gibbon accumulated [^3^H]-estradiol-17β-glucuronide and [^3^H]-folic acid, but not [^3^H]-estrone sulfate and [^3^H]-taurocholic acid, similar to the SLC22A10 isoform with 552 amino acids (***Fig. 4C***, ***Supplementary Fig. 4***). Moreover, we observed that longer isoforms of SLC22A10 created for the bonobo and orangutan, which were not predicted to express these isoforms (552 amino acids), accumulated [^3^H]-estradiol-17β-glucuronide, and were also expressed on the plasma membrane (***Supplementary Fig. 4***).

**Figure 4.**
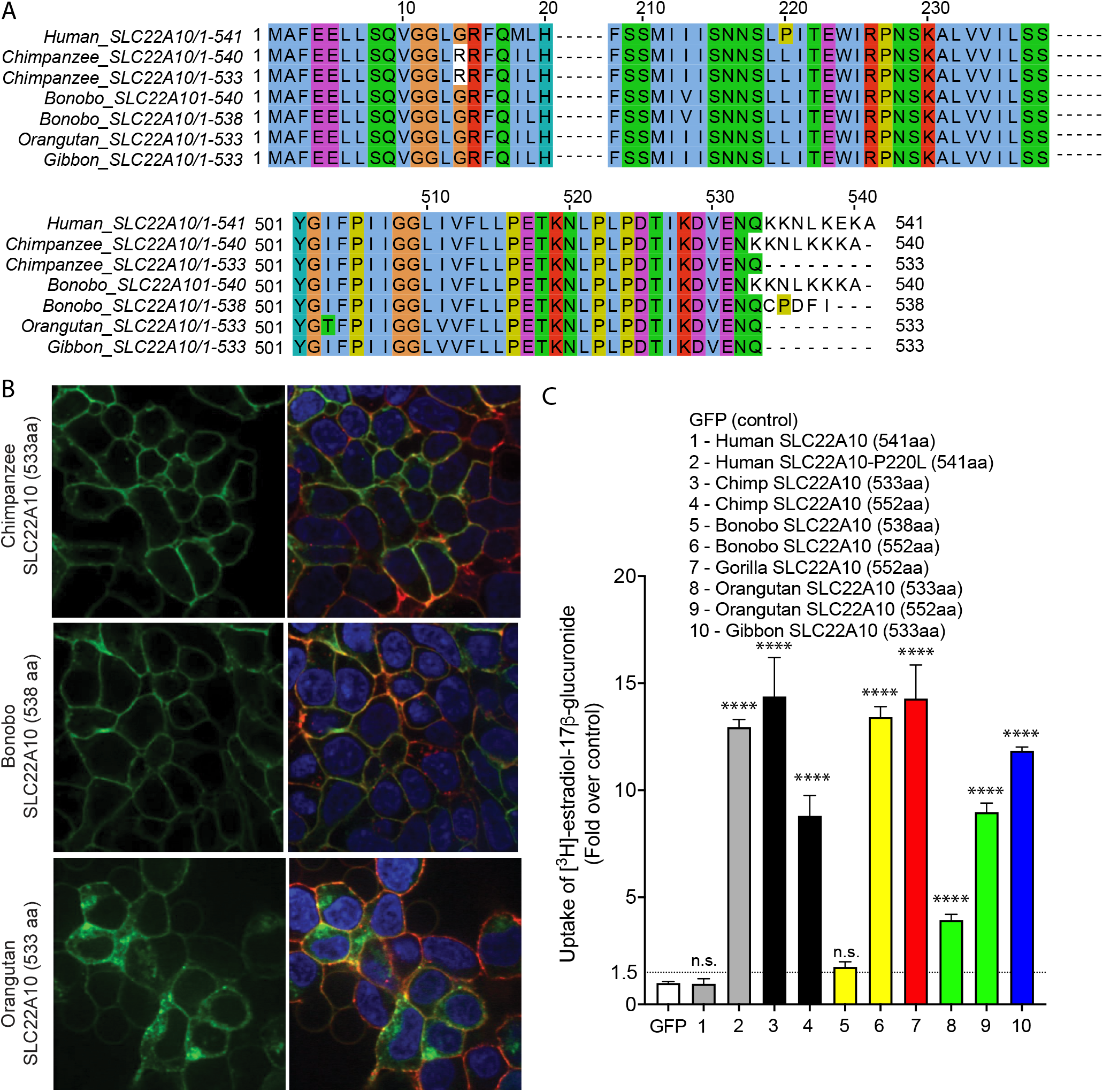
SLC22A10 of chimpanzees, bonobos, orangutans, and gibbons are predicted to have shorter isoforms expressing 533 or 538 amino acids. **A**. A comparison of the SLC22A10 amino acid sequence of humans, chimpanzees, bonobos, and orangutans, which express 533 (chimpanzee, orangutan, gibbon), 538 (bonobo), 540 (bonobo, chimpanzee), or 541 (human) amino acids, shows that the major differences are at the end of the SLC22A10 sequence. **B**. Confocal imaging revealed that SLC22A10 from chimpanzees and bonobos (isoforms expressing 533 or 538 amino acids) primarily localize to the plasma membrane of the cell, whereas weaker localization was observed for orangutan SLC22A10 (533 amino acids) to the plasma membrane of the cell. GFP conjugated to SLC22A10 was used for this experiment. **D.** The uptake of [^3^H]-estradiol-17b-glucuronide in HEK293 cells was observed after transient transfection of plasmids encoding human SLC22A10 with reference amino acids or SLC22A10 with reference amino acids of other great apes with different isoforms. The results showed that SLC22A10 isoforms expressing 533 and 552 amino acids significantly accumulate the substrate. However, weaker substrate accumulation was observed in cells transfected with the bonobo SLC22A10 isoform expressing 538 amino acids.

### Uptake of various steroid glucuronides by chimpanzee SLC22A10 reveals that 17β-glucuronides are preferred over 3α-glucuronide conjugates

Our experiments revealed significant accumulation of estradiol-3β-glucuronide, estradiol-17β-glucuronide, estrone-3β-glucuronide and testosterone-17β-glucuronide (***Fig. 5A***). We also observed weak but significant accumulation of androstanediol-3α-glucuronide, but no significant accumulation of androsterone-3-glucuronide, etiocholanolone-3α-glucuronide, progesterone, and testosterone (***Fig. 5A***). Although the data are limited, it is intriguing to note the substrate preference of SLC22A10. Specifically, 17β-glucuronides appear to be favored over 3α-glucuronide conjugates.

**Figure 5.**
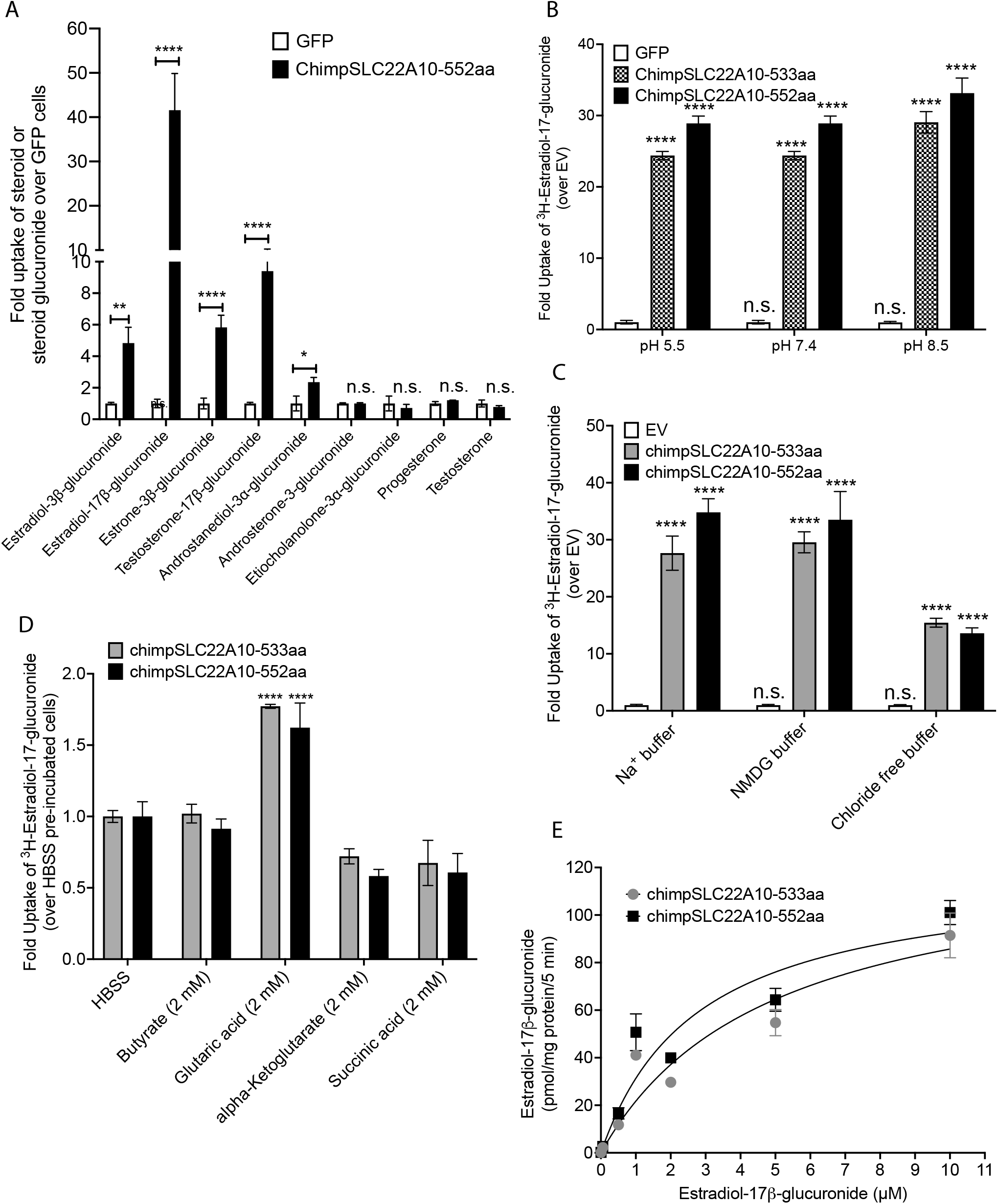
This figure presents information about the transport mechanism and kinetics of chimpanzee SLC22A10. **A.** The uptake of seven steroid glucuronides and two steroids in HEK293 cells stably transfected with chimpanzee SLC22A10 (552 amino acids) was measured using LC/MS-MS to determine the accumulation of the compounds. **B.** The uptake of seven steroid glucuronides and two steroids in HEK293 cells stably transfected with chimpanzee SLC22A10 (552 amino acids) was measured using LC/MS-MS to determine the accumulation of the compounds. **C.** The effect of pH on accumulation of [^3^H]-estradiol-17b-glucuronide in HEK293 cells stably transfected with chimpanzee SLC22A10 isoforms expressing 533 and 552 amino acids was investigated. **D.** The effect of sodium and chloride on accumulation of [^3^H]-estradiol-17b-glucuronide in HEK293 cells stably transfected with 533 and 552 amino acid isoforms of chimpanzee SLC22A10 was investigated. **E.** The effects of *trans*-stimulation of [^3^H]-estradiol-17b-glucuronide uptake by chimpanzee SLC22A10 was determined. Uptake was *trans*-stimulated by preloading the cells with 2 mM of butyrate, glutaric acid, alpha-ketoglutarate, or succinic acid for 2 hours, and then measuring the uptake of [^3^H]-estradiol-17b-glucuronide after 15 minutes. The data are presented as mean ± S.D. and were normalized by setting the uptake of SLC22A10-expressing cells trans-stimulated by HBSS to 1.0. *Trans*-stimulation of [^3^H]-estradiol-17b-glucuronide by glutaric acid was observed for both isoforms of chimpanzee SLC22A10. **E**. The kinetics of [^3^H]-estradiol-17b-glucuronide uptake for chimpanzee SLC22A10 isoforms expressing 533 and 552 amino acids were analyzed. The uptake rate was evaluated at 5 minutes and the data were fit to a Michaelis-Menten equation. To fit the kinetic curve to a Michaelis-Menten equation, the concentration of estradiol-17b-glucuronide is set up to 10 µM. The figure shows a representative plot from one experiment. All experiments were repeated once, in triplicate and showed similar results.

### Chimpanzee SLC22A10-mediated uptake is sodium independent, pH independent, chloride independent and trans-stimulated by glutaric acid

Transporters in the SLC22 family may be secondary active, in which case they rely on various sources of energy to mediate active flux of their substrates. Accordingly, we examined the transport mechanism for two isoforms (533 and 552 amino acids) of chimpanzee SLC22A10. At the three pH levels evaluated, the uptake of [^3^H]-estradiol-17β-glucuronide by both isoforms of chimpanzee SLC22A10 were similar (***Fig. 5B***). While the uptake was not dependent on sodium (***Fig. 5C***), it was significantly reduced in the absence of chloride in the buffer (***Fig. 5C***). Several organic anion transporters in the SLC22 family are *trans*-stimulated by dicarboxylic acids^3, 16^. Similarly, we observed that the uptake of [^3^H]-estradiol-17β-glucuronide was *trans*-stimulated by glutaric acid but not significantly by other dicarboxylates (α-ketoglutarate and succinic acid) or the monocarboxylic acid, butyrate (***Fig. 5D***). The uptake kinetics of [^3^H]-estradiol-17β-glucuronide exhibited saturable characteristics with virtually identical Km values for the two protein isoforms at 3.28 ± 2.25 µM and 2.16 ± 0.59 µM for the 533 aa and 552 aa isoforms, respectively (***Fig. 5E***).

## Discussion

Transporters play a major role in total body homeostasis as they function to regulate the levels of many solutes including endogenous metabolites, essential nutrients, and environmental toxins. Over 30% of genes in the Solute Carrier superfamily have no known function^1^. Identifying the substrates or deorphaning transporters presents significant challenges due to their diverse structural and functional characteristics, as well as the intricate cellular and physiological context in which they operate.

Overcoming these challenges necessitates the implementation of innovative approaches that integrate computational predictions, high-throughput screening, functional assays, and targeted experimental investigations^1, 17^. Recent advancements in transporter research in the past four years have led to the identification of ligands for several transporters in the organic ion transporter family, SLC22. Notably, SLC22A14, SLC22A15, and SLC22A24 have been successfully characterized^2, 3, 18^. In this study, we pursued a distinct approach to deorphaning a human SLC22 family member, SLC22A10 by investigating the function of great ape orthologs. Our comprehensive investigations have yielded five major findings that contribute to our understanding of the function and evolution of SLC22A10 in higher order primates.

First, SLC22A10 functions as a steroid glucuronide transporter in great apes (as shown in ***Fig. 1*** and ***Fig. 2***). Unlike other organic anion transporters in the SLC22 family such as SLC22A6, SLC22A7, and SLC22A8, which transport a variety of organic anions including uric acid, steroid glucuronides, bile acids, steroid sulfates, dicarboxylic acid, and phosphate-containing nucleotides^6^, the ortholog of SLC22A10 from primates (chimpanzee, bonobo, gorilla, orangutan and gibbon) transported primarily estradiol-17β-glucuronide (***Fig. S1***) with significantly weaker uptake of other organic anions such as folic acid and methotrexate (***Fig. S1*** and ***Fig. S4***). Interestingly, the SLC22A10 ortholog in the squirrel monkey, a New World monkey, preferred estrone sulfate over estradiol-17β-glucuronide (***Fig. S5***) whereas the gene encoding SLC22A10 is absent in Old World monkeys (*Cercopithecidae* family) (ensemble release 110). These results suggest an evolving role of the function of the transporter in non-human primates. In both chimpanzee and squirrel monkey, SLC22A10 is expressed specifically in the liver (NHPRTR, http://nhprtr.org/), consistent with a role in steroid metabolism. The closest paralog of SLC22A10 is SLC22A24, a recently deorphaned transporter^3^. Both transporters take up steroid conjugates though chimpanzee SLC22A22A10 has a narrower substrate specificity than the human SLC22A24.

Our second major finding is that the function of SLC22A10 is lost in humans. That is, while human SLC22A10 is transcribed, the protein expression is undetectable in human cell lines transfected with human SLC22A10 and in human liver tissue^19–23^ (https://www.proteinatlas.org/), consistent with a loss of function of the gene in humans. Indeed, proteomic analysis demonstrated that the reference proline (Pro220) at the 220 amino acid position of human SLC22A10 displayed significantly lower protein expression compared to the mutated form, Leu220 (see **Table 1**). Further, as Pro220 is fixed in humans resulting in a loss of function, the gene has been under less selective pressure. Not surprisingly, SLC22A10 harbors a prevalent nonsense variant, p.Trp96Ter (rs1790218), which is frequently observed at high allele frequencies in human populations ranging from ∼20% in African to 50% in European (***Fig. S5***). Nonsense-mediated decay (NMD) is known to be triggered by premature stop codons. The Genotype-Tissue Expression (GTEx) project uncovered a significant association between the rs1790218 variant, which encodes the A-allele and p.Trp96Ter, and a considerable decrease in the transcript level of SLC22A10 samples (source: https://gtexportal.org/home/snp/rs1790218). This finding suggests that individuals carrying one copy of the p.Trp96Ter are likely to exhibit an even lower protein signal in the liver in comparison to individuals that harbor p.Trp96. This finding is relevant to how the protein itself (though not a functioning protein) is being lost. During our efforts to clone the human SLC22A10 gene using pooled human liver samples, we selected 27 individual colonies for further analysis and subsequent sequencing. Only four of these colonies contained the complete transcript spanning 1626 base pairs. The remaining 23 out of 27 colonies selected had a identical 219 bp deletion, resulting in a truncated sequence of 1407 bp, corresponding to 469 amino acids (***Fig. S6***). However, NCBI predicted a different deletion of 324 bp, starting at the same GT splice donor site that we observed, but extending to a further GT acceptor site (NCBI Reference Sequence: XM_047426921).

Our third major finding is that Leu220Pro alone caused the loss of function of the gene in hominins. That is, a single proline substitution resulted in no expression of SLC22A10 on the plasma membrane and significantly reduced protein abundance (**Table1**). When Pro220 in human SLC22A10 was mutated to Leu220, it acquired the functional capacity observed in chimpanzee SLC22A10, resulting in substantial accumulation of the substrate, estradiol-17β-glucuronide. Conversely, when the amino acid at position 220 in chimpanzee SLC22A10 was mutated to proline, the uptake of estradiol-17β-glucuronide was completely abolished (***Fig. 3B***). Cordes F. et al. (2002) reported that distortions of transmembrane helices can be induced by the presence of proline^24^. Using AlphaFold2 model of human SLC22A10, we observed that the proline 220 in humans introduces a kink in the alpha-helix, which in turn can affect the conformation of the 225-230 loop and, consequently increasing the accessibility of the lysine 230 for ubiquitination. In addition, based on the ΔΔG calculation the proline 220 is significantly less stable with a higher ΔΔG. Based on comprehensive genomics datasets from large-scale populations such as gnomAD^25^, GDBIG (http://www.bigcs.com.cn/), TopMed Freeze 8^26^, and GME Variome^27^, there have been no reported individuals with SLC22A10-Leu220. Additionally, four high-coverage archaic hominin genomes – three Neanderthals and a Denisovan are homozygous for the C allele (chr11:63297455 (hg38)), suggesting that this mutation emerged following Pan-Homo divergence and before modern humans diverged from archaic hominins. While the SLC22A10-Trp96Ter variant has not been assayed in archaic hominin or ancient human genomes, two SNPs in strong LD with this variant occur in ancient human genomes at least 30,000 years ago (***Fig. S7***) and the variant itself is estimated to emerge ∼120,000 years ago (***Fig. S8***).

Our fourth finding unveils the transporter activity and expression of the predicted isoforms of SLC22A10 on the plasma membrane in great apes (refer to ***Fig. 2***, ***Fig. 3***, and ***Fig. 4***). Importantly, among the great apes, three isoforms were initially predicted: 533, 540, and 552 amino acids in length. However, further investigation revealed that only the isoforms with 533 and 552 amino acids were actually expressed on the plasma membrane and exhibited transporter activity, as depicted in **Figure 4** and **Figure 5**. In **Figure 2C**, the last exons (exon number 10) for humans and other great apes is presented. It is worth noting that human SLC22A10 is predicted to consist of 541 amino acids. The gene annotation of chimpanzees and other primates in Ensembl is limited by an anthropocentric approach that heavily relies on human annotation as a reference. In the current version 110 of Ensemble.org, the chimpanzee, gorilla, bonobo gene are annotated with 540 amino acids. This annotation utilizes a non-canonical splice site at the end of exon 9 (‘CA’) instead of the canonical donor splice site (’GT’), which is not supported by our chimpanzee liver RNA-seq data^12^ (***Fig. S3*** and ***Fig. 2C***). In contrast, NCBI predicted that chimpanzee possess isoforms with 533 and 552 amino acids, which is consistent with our observation from chimpanzee liver RNAseq data^12^. Within the SLC22A family, several transporters, including SLC22A7^28^ and SLC22A24^3^ exhibit distinct isoforms.

Overall, our studies revealed that the activity of SLC22A10 has evolved in primates, with Old World monkeys lack SLC22A10 orthologs, and New World monkeys exhibit a different substrate preference compared to great apes. In addition, the great apes SLC22A10 was rendered nonfunctional by a single missense mutation during hominin evolution after our shared ancestor with the chimpanzee. This missense mutation resulted in a complete loss of human SLC22A10 transporter activity, due to lack of protein expression on plasma membrane and reduce protein abundance. With time, the gene has accumulated additional mutations, including a stop codon (p.Trp96Ter), which has led to reductions in the levels of the mRNA transcript and corresponding reductions in protein level. This gene exhibits features that classify it as unitary pseudogene^29, 30^. A pseudogene can be defined by the loss of the original function due to errors during transcription or translation, or as a gene producing a protein that does not have the same functional repertoire as the original gene^30^. Consequently, a pseudogene will not necessarily evolve under a neutral theory of molecular evolution. Pseudogenes can be categorized into 3 different types depending on their functional state. These include exapted pseudogenes, which have gained a new biological function; “dying” pseudogenes, which still have some transcriptional activity; and “dead” pseudogenes, which do not exhibit any signs of activity and evolve under the neutral theory^29^. Based on the evidence at hand, we cannot differentiate if the pseudogene is exapted or dying. However, the SLC22A10 gene is a well-established gene that originated from the last common ancestor of boreoeutheria, with no functioning counterparts in the human genome. Thus, we can classify human SLC22A10 an unitary pseudogene. Examples of unitary pseudogene in human is MUP (major urinary protein), whereas uricase (UOX) and GULO (L-Gulonolactone oxidase) are well established unitary pseudogenes that were inactivated before the separation of human and chimpanzee^31^, There are known pseudogenes in the SLC superfamily (SLC22A20, SLC35E2A, SLC6A10, SLC23A4P, SLC6A21P)^32^, however none of them are due to a single missense mutation. Future studies are needed to determine whether the loss of function human SLC22A10-P220 is a favorable situation for humans and whether similar mechanisms have led to the inactivation of other orphan genes in the human genome.

## Materials and Methods

### Generation of SLC22A10 ortholog cDNA constructs

Sequences of SLC22A10 orthologs from various species, along with their respective sequence IDs, were obtained from ensembl.org (release version 104)^33^ and the National Center for Biotechnology Information (NCBI). The gene’s cDNA was synthesized using GenScript Gene Synthesis and DNA Synthesis Services and subsequently inserted into the multiple cloning sites on the expression vectors pcDNA3.1(+), pcDNA3.1(+)C-eGFP, or pcDNA3.1(+)N-eGFP. The sequence of the constructs was verified through sequencing conducted by GenScript.

Human SLC22A10, NM_001039752, NP_001034841.3 (541 aa)

Chimpanzee SLC22A10, XM_016921102.2, XP_016776591.1 (552 aa)

Chimpanzee SLC22A10, ENSPTRT00000087785.1, A0A2I3TWY5 (540 aa)

Chimpanzee SLC22A10, XM_024347215.1, XP_024202983 (533 aa)

Bonobo SLC22A10, XM_034932815.1, XP_034788706 (538 aa)

Bonobo SLC22A10, ENSPPAT00000058172.1, A0A2R9CA90 (540 aa)

Gorilla SLC22A10, XM_019037494.1, XP_018893039 (552 aa)

Gorilla SLC22A10, ENSGGOT00000014974.3, G3RG24 (540 aa)

Orangutan SLC22A10, XM_024255628.1, XP_024111396.1 (533 aa)

Orangutan SLC22A10, XM_054439153.1, XP_054295128.1 (552 aa)

Gibbon SLC22A10, XM_003274123.2, XP_003274171.2 (533 aa)

Squirrel monkey SLC22A10, ENSSBOT00000021267.1, A0A2K6SBP5 (552 aa)

### Site-directed mutagenesis

Site-directed mutagenesis to create Methionine18 to Isoleucine18 in human SLC22A10, Proline220 to Leucine220 or vice versa in chimpanzee SLC22A10 and human SLC22A10, were performed using Q5 Site-Directed Mutagenesis Protocol from NEB (#E0554). NEBaseChanger, https://nebasechanger.neb.com/, is used to assist in the design of primers for the site-directed mutagenesis experiment.

Primers for p.M18I:

mut_hsA10_c54C_F: GATTTCAGATcCTTCATCTGGTTTTTATTCTTC

mut_hsA10_c54C_R: TCCCAAGGCCTCCAACTT

Primers for p.P220L:

mut_hsA10_c659T_F: AATTCTTTGCtCATTACTGAG

mut_hsA10_c659T_R: ATTTGATATAATGATCATGGAAG

Primers for p.L220P:

mut_ptA10_c659C_F: AATTCTTTGCcCATTACTGAGTG

mut_ptA10_c659C_R: ATTTGATATAATGATCATGGAAG

Primers for deletion of A nucleotide in position c1607 and then inserting the tail at the last exon for human to chimpanzee (consist of 34 bp):

mut_hsA10_c1607Adel_F: TCTCAAGGAAAAGGCATAAATG

mut_hsA10_c1607Adel_R: TTTTTTTTTGATTTTCCACATCC

mut_hsA10_c1622ins_F: caaaagtgaccaaatttTAACTCGAGTCTAGAGGG

mut_hsA10_c1622ins_R: tgtagcaatcatttttaTGCCTTTTCCTTGAGATTTTTTTTTG

### Cell culture

Human embryonic kidney cell lines (HEK293) containing a Flp-In expression vector (HEK293 Flp-In) obtained from ThermoFisher Scientific were utilized in this study. These cells have previously been employed for the generation of stable cell lines for transporter assays^2, 3, 16, 28^. The HEK293-FlpIn cells from ATCC were cultured in DMEM, high glucose (#11965118, ThermoFisher Scientific) supplemented with 10% fetal bovine serum (heat inactivate, #10438026, ThermoFisher Scientific). Penicillin-Streptomycin (#15070063, ThermoFisher Scientific) was added to DMEM media (50 unit/500 mL DMEM). During transfection and when cells were plated for transporter studies, media without penicillin/streptomycin supplementation were used. The cells were regularly screened for mycoplasma contamination (MycoProbe Mycoplasma Detection Kit, #CUL001B, Fisher).

### Generation of cells transiently or stably expressing cDNAs

Expression vectors of SLC22A10 orthologs were introduced into HEK293 Flp-In cells either through transient transfection or stable transfection using Lipofectamine LTX (Thermo Fisher Scientific). For transfections in a 48-well plate (seeding density: 1.0×10^5^ cells/well), 200 ng of DNA and 0.4 μL of Lipofectamine LTX were utilized, while for transfections in a 100 mm tissue culture plate (seeding density: 4×10^6^ cells/well), 10 μg of DNA and 44 μL of Lipofectamine LTX were used. More comprehensive methods for generating transiently or stably transfected cells have been described in our previous work (see reference^2, 3^). In the case of transient transfection, cells were used for transporter studies (refer to the section titled “Transporter uptake studies“) after 36-48 hours or for protein quantification after 72 hours. To establish stable cell lines, 3000 ng of DNA (SLC22A10 ortholog expression vectors) and 10.5 μL of Lipofectamine LTX were employed to transfect HEK293 Flp-In cells seeded in a 6-well plate (seeding density: 7-8×10^5^ cells/well). After 48 hours, cells were transferred to a new 100 mm tissue culture plate and treated with 800 μg/mL Geneticin. Fresh media containing 800 μg/mL Geneticin was replenished every other day for 1 week. Stable cell lines were utilized for confocal imaging to determine the plasma membrane localization of SLC22A10 orthologs and their various isoforms. Unless specified otherwise, stable cell lines were used for transporter assays.

### Fluorescence microscopy

For the immunostaining experiments, HEK293 Flp-In stable cell lines expressing different SLC22A10 orthologs were cultured on poly-D-lysine-treated 12-well plates with sterile coverslips at a density of 200,000 cells per well. After two days of seeding when the cells reached 90-100% confluency, the staining procedure was conducted. On the day of staining, the cell culture media was carefully removed, and the cells were washed using cold Hank’s Balanced Salt Solution (HBSS, #14025092, ThermoFisher Scientific). To initiate the staining process, the plasma membrane was first labeled using Wheat Germ Agglutin (WGA) Alexa Fluor 647 conjugate (Invitrogen Life Sciences Corporation) diluted at a ratio of 1:500 in HBSS, followed by a 15-minute incubation at room temperature. Following the staining step, the WGA solution was aspirated, and the cells were washed three times with HBSS. Subsequently, the cells were fixed with a solution of 3.7% formaldehyde in HBSS for 20 minutes. After the fixation step, the cells were washed three times with HBSS. To stain the nucleus, Hoechst solution (ThermoFisher Scientific Inc.) diluted at a ratio of 1:2000 in HBSS was applied to the cells and incubated for 20 minutes at RT, in darkness. After the staining period, the Hoechst solution was aspirated, and the cells were washed twice with HBSS. The coverslips were carefully mounted on Superfrost Plus Microscope Slides (ThermoFisher Scientific) using a small amount of SlowFade^TM^ Gold Antifade Mountant (#S36940, ThermoFisher Scientific). The mounted slides were left to dry overnight in darkness before being imaged using an inverted Nikon Ti microscope equipped with a CSU-22 spinning disk confocal system available at the Center for Advanced Light Microscopy (CALM) at University of California San Franciso. The image acquisition settings were as follows: DAPI channel with a 300ms exposure time and 50% laser power, FITC channel with a 300ms exposure time and 25% laser power, and CY5 channel with a 100ms exposure time and 5% laser power. Image alignment and merging were performed using Fiji software. This experimental protocol has been previously utilized and described in our published work^34, 35^.

### Transporter uptake studies

HEK293 Flp-In cells expressing SLC22A10 were seeded at a density of 120,000 to 150,000 cells/0.3 mL in poly-D-lysine-coated 48-well plates approximately 16 to 24 hours prior to conducting uptake studies. The uptake studies for transporters, as detailed below, are methods we have previously described^2, 3^. For transiently expressing SLC22A10, the methods outlined in the previous section pertaining to transient expression in HEK293 Flp-In cells were followed prior to this step. Prior to uptake studies, the culture medium (Dulbecco’s modified Eagle’s medium, DMEM) supplemented with 10% fetal bovine serum was aspirated, and the cells were incubated in 0.8 mL of Hank’s Balanced Salt Solution (HBSS) at 37 °C for 10-20 minutes. For screening radiolabeled compounds as SLC22A10 substrates, minute quantities of radiolabeled compounds (^3^H or ^14^C) were diluted in HBSS (at ratios of 1:2000 or 1:3000) for uptake experiments. Unlabeled compounds were added to obtain specific concentrations, which are described in the Results section or figure legends along with the uptake times. Uptake reactions were terminated by washing the cells twice with 0.8 mL of HBSS buffer, followed by incubation in 750 μL of lysis buffer (0.1 N NaOH, 0.1% v/v SDS). A 690 μL portion of the cell lysate was transferred to scintillation fluid for scintillation counting. For pH dependence experiments, the HBSS buffer was adjusted to different pH levels (5.5, 7.4, and 8.5) using hydrochloric acid or sodium hydroxide. For sodium and chloride dependence studies, three distinct uptake buffers were employed: (1) chloride-free buffer (composed of 125 mM sodium gluconate, 4.8 mM potassium gluconate, 1.2 mM magnesium sulfate, 1.3 mM calcium gluconate, and 5 mM HEPES; adjusted to pH 7.4 with sodium hydroxide); or (2) sodium buffer (composed of 140 mM sodium chloride, 4.73 mM potassium chloride, 1.25 mM calcium chloride, 1.25 mM magnesium sulfate, and 5 mM HEPES, adjusted to pH 7.4 with sodium hydroxide); or (3) sodium-free buffer (composed of 140 mM N-methyl-D-glucamine chloride, 1.25 mM magnesium sulfate, 4.73 mM potassium chloride and 1.25 mM calcium chloride), adjusted to pH 7.4 with potassium hydroxide). For trans-stimulation studies, the experimental conditions described in our previously published methods were followed^2, 3^. In brief, the SLC22A10 or EV stable cell lines were pre-incubated with either buffer or 2 mM succinic acid, 2 mM α-ketoglutaric acid, 2 mM butyric acid, or 2 mM glutaric acid for 2 hours. Subsequently, the cells were washed twice with HBSS before commencing the uptake of the anions (estradiol-17β-glucuronide).

### Kinetic studies of estradiol glucuronide

Kinetic studies of estradiol-17β-glucuronide were conducted in HEK293 Flp-In cells expressing chimpanzee SLC22A10 isoforms (533 amino acid and 552 amino acid) that were stably transfected. The experimental conditions for the kinetic studies closely followed the methods previously published by our research group (reference provided). Initially, we examined the time-dependent uptake of the substrates using trace amounts of the radioactive compound. Concentrations of the non-labeled compounds were varied up to 50 µM. For the kinetic studies, a duration of five minutes at 37°C was chosen as it fell within the linear range observed in the uptake versus time plot for each substrate. Each data point represents the mean ± standard deviation of uptake in the cells transfected with the transporter, subtracted by that in empty vector cells. The obtained data were fitted to a Michaelis-Menten equation to estimate the kinetic parameters. Plots were generated based on a representative experiment out of three independent studies.

### Protein extraction and global proteomics of HEK293 cells expressing SLC22A10 orthologs

HEK293 Flp-In cells were transfected transiently with various SLC22A10 orthologs, including human, chimpanzee, and the mutations to proline or leucine at position 220. After 72 hours of transfection, cell pellets were collected and shipped to Dr. Per Artursson’s laboratory in Uppsala University for protein quantification. The quantification was performed on both HEK293 cells and HEK293 cells expressing the different SLC22A10 orthologs and mutations. HEK293 cell pellets (50–92 mg) were lysed in a lysis buffer containing 50 mM dithiothreitol, 2% sodium dodecyl sulfate in 100 mM Tris/HCl pH 7.8. The lysates were incubated at 95°C for 5 min and sonicated with 20 pulses of 1 second, 20% amplitude by using a sonicator coupled with a microtip probe. The lysates were centrifuged at 14,000×g for 10 min and supernatants were collected. Using LysC and trypsin, the multi-enzyme digestion filter-aided sample preparation (MED-FASP) approach was performed^36^. C18 stage tips were used to desalt the peptide mixture^37, 38^ and samples were stored at -20°C until analysis. Protein and peptide content were determined by using tryptophan fluorescence assay^39^. The global proteomics analysis was performed on a Q Exactive HF mass spectrometer (Thermo Fisher Scientific) coupled to a nano–liquid chromatography (nLC). EASY-spray C18-column (50 cm long, 75 µm inner diameter) was used to separate peptides on a ACN/water gradient (with 0.1% formic acid) over 150 min. MS was set to data dependent acquisition with a Top-N method (full MS followed by ddMS2 scans). Proteins were identified using MaxQuant software (version 2.1.0.0)^40^ with the human proteome reference from UniProtKB (October, 2022). Total protein approach was used as the protein quantification method^41^.

### RNA isolation and quantitative RT-PCR

HEK293 Flp-In cells were cultured in poly-D-lysine coated 24-well plates at a seeding density of 1.5-1.8 × 10^5^ cells per well, allowing them to reach 75-80% confluency. The RT-PCR method for transcript levels determination as detailed bleow, are methods we have previously described^3^. Once the desired confluency was achieved, the cells were transiently transfected with either the vector alone or the vector containing different SLC22A10 orthologs (in the pcDNA3.1(+) expression vector). For the transfection mixture, 500 ng of plasmid DNA, 1 μL of Lipofectamine LTX (Thermo Fisher Scientific), and 100 μL of Opti-MEM I reduced serum media (Thermo Fisher Scientific) were used. After 36-48 hours of transfection, the media was removed, and RNA Lysis buffer (350 μL) was added to each well. Total RNA was isolated from the cells using the Qiagen RNeasy kit (Qiagen). Subsequently, cDNA was synthesized using the SuperScript VILO cDNA Synthesis Kit (ThermoFisher Scientific). For quantitative RT-PCR (qRT-PCR), Taqman reagents and specific primer and probe sets were used, targeting human SLC22A10 (Assay ID: Hs01397962_m1) and beta actin (Assay ID: Hs99999903_m1) (Applied Biosystems, Foster City, CA). The qRT-PCR reactions were conducted in a 96-well plate, with a reaction volume of 10 μL, using the QuantStudio™ 6 Flex Real-Time PCR System and the default instrument settings. The expression levels were determined using the Ct method, and the data were normalized to the endogenous levels of beta actin. The results are presented as fold-increases in the SLC22A10 transcript levels relative to the cell lines expressing the vector control. The analysis was based on three independent biological samples.

### Cloning of SLC22A10 in pooled human liver

For the cloning process, pooled total RNA samples from human liver were obtained from Clontech. Each sample (2 μg) of total RNA was reverse transcribed into cDNA using the SuperScript VILO cDNA Synthesis kit (Thermo Fisher Scientific) following the manufacturer’s instructions. The primers specified below were employed for PCR amplification of the NM_001039752 transcript: Forward primer: ACCGAGCTCGGATCC<u>ATGGCCTTTGAGGAGCTC</u>; and reverse primer: CCCTCTAGACTCGAG<u>TTATGCCTTTTCCTTGAGATT</u>. The nucleotide underlined are open reading frame of SLC22A10. The resulting PCR products were cloned into BamHI and XhoI multiple-cloning site of the pcDNA5FRT vector and subsequently subjected to sequencing at MCLAB in South San Francisco to determine the sequence of the transcript. For the cloning of human SLC22A10, the KOD Xtreme Hot Start DNA polymerase kit (Takara) was utilized. The PCR cycling conditions were as follows: (i) initial activation at 94°C for 2 minutes, (ii) denaturation at 98°C for 10 seconds, (iii) annealing at 57.5°C for 30 seconds, and (iv) extension at 68°C for 1 minute.

### Calculating ΔΔG with the PRISM’s rosetta_ddG_pipelne v 0.2.4^14^ using Rosetta v 3.15

In brief, the full-length human SLC22A10-P220 (Uniprot ID Q63ZE4) structure predicted by the AlphaFold^42^ from the AlphaFold DB^43^ was oriented in the membrane with PPM 3.0 web server^44^ with the default settings and relaxed using the Rosetta’s relax protocol^44^. The best of 20 generated structures was used for the ΔΔG calculations with the cartesian_ddG protocol^45^ repeated in 5 replicas. ΔΔE scores were calculated using GEMME^46^ and rank-normalized as in the reference paper^14^. The relaxed structure, ΔΔG, and ΔΔE results are available at zenodo.

### Transporter uptake studies and LC/MS/MS analysis

The list of steroid conjugates and their sources can be found in the ***Table S1***. Each steroid conjugates was dissolved in DMSO to obtain 20 mM stock solution. Compounds were stored in -20°C freezer.

HEK293 Flp-In cells stably transfected with GFP only, chimpanzee SLC22A10 (533 amino acid) and chimpanzee SLC22A10 (552 amino acid) were plated in poly-D-lysine coated 48-well plates at a seeding density of 1.5 × 10^5^ cells per well, allowing them to reach 90-95% confluency after 16-24 hours. Prior to uptake studies, the culture medium (Dulbecco’s modified Eagle’s medium, DMEM) supplemented with 10% fetal bovine serum was aspirated, and the cells were incubated in 0.8 mL of Hank’s Balanced Salt Solution (HBSS) at 37 °C for 10-20 minutes. To screen various steroid and steroid conjugate compounds as SLC22A10 substrates, HEK293 Flp-In cells stably expressing GFP or chimpanzee SLC22A10 were incubated with HBSS buffer containing 10 µM of the respective compounds for 20 minutes. The uptake reactions were terminated by washing the cells twice with 0.8 mL of HBSS buffer, followed by incubation in 400 μL of methanol. After 30 minutes of shaking at room temperature, 300 µL of methanol containing the extracted steroid or steroid conjugates from each well were transferred to a 1.5 mL tube and stored at -80°C before quantification using LC/MS/MS analysis.

Subaliquot of cellular extracts (90 µL) were spiked with 10 µL of deuterated 17β-estradiol glucuronide, mixed by vortexing, and filtered at 0.2µm through polyvinyl difluoride membranes (Agilent Technologies, Santa Rosa, CA, USA) by centrifugation and 10,000g. After filtration, samples were enriched with 25nM 1-cyclohexyl-3-uriedo-decanoic acid (Sigma-Adlrich, St. Lousi MO) as an internal standard. Metabolites were measured using ultra-performance liquid chromatography–electrospray ionization tandem mass spectrometry (UPLC-ESI-MS/MS) on a API 4500 QQQ (Sciex, Framingham, MA) with a scheduled multiple reaction monitoring (MRM) using methods adapted from Ke et.al, 2015^47^. Analytes were separated on a Waters I-Class UPLC-FTN equipped with a 2.1 × 100 mm i.d., 1.7 μm Acquity BEH C_18_ column (Waters Co; Milford, MA) held at 50°C. Analytes in 5µL injections were separated using water (solvent A) and methanol (solvent B) both containing 2 mM ammonium formate at 400 µL/min with the following gradient: Initial 40% B to 70% B at 2 min, to 98%B at 3 min, held to 4 min, to 40%B at 5.1 min held to 6 min. Mass spectrometer acquisition parameters and analyte retention times are described in ***Table S2***.

### Sequencing data processing and analyses of SLC22A10 in greater apes

Orthologous genome regions of *SLC22A10* coding sequence in multiple primate species were obtained using UCSC liftOver (default parameters) (Hinrichs et al. 2006) based on hg38 (human), panTro6 (chimpanzee), panPan3 (bonobo), gorGor6 (gorilla) and ponAbe3 (orangutan) assemblies. These regions were aligned using MUSCLE (Edgar 2004) and visualized using MView (Brown, Leroy, and Sander 1998). *SLC22A10* exons and coding sequences are based on RefSeq annotations (O’Leary et al. 2016) for human, chimpanzee and gorilla. Despite the long *SLC22A10* isoform not being annotated in RefSeq for bonobo and orangutan, we included these two species in the alignment considering the plausibility of the protein model in the context of their genomes and the high genomic conservation in comparison to chimpanzee and gorilla.

Allele frequencies from non-human great apes were obtained from available whole-genome sequencing data including 59 chimpanzees, 10 bonobos, 49 gorillas and 16 orangutans^48–51^. All samples were mapped to Hg19. We extracted the genotyping information in position chr11:63078478 (Hg19 coordinates) to calculate the allele frequencies per population. We manually curated the genotypes by checking the raw reads overlapping this region in the BAM files. In chimpanzees we report a frequency of the insertion to be 3/108=2.5%; in bonobos is 0/20=0%; in gorillas is 0/98=0%; and in orangutans is 0/32=0%. The global human allele frequencies were obtained from 1000 Genomes Project Phase 3^52^ database in Ensembl for the rs562147200 SNP.

### SLC22A10 variants in archaic hominins and ancient humans

Archaic hominin genotypes for three Neanderthals and a Denisovan were retrieved from http://ftp.eva.mpg.de/neandertal/Vindija/VCF/ and http://ftp.eva.mpg.de/neandertal/Chagyrskaya/VCF/^53–56^. We used the LDproxy tool from LDlink^57^ to identify variants in high LD with rs1790218 in all Thousand Genomes populations and intersected these variants with those assayed in ancient humans from the Allen Ancient DNA Resource (AADR)^58^, retrieved from https://reichdata.hms.harvard.edu/pub/datasets/amh_repo/curated_releases/V54/V54.1.p1/SHARE/public. dir/v54.1.p1_1240K_public.tar. We identified two such variants: rs1783634 (D’ = 1, r2 = 0.9839) and rs1201559 (D’ = 0.9975, r2 = 0.9775). We filtered the AADR genotypes for these SNPs, excluding samples from archaic hominins and individuals that were not genotyped at both loci. We calculated allele frequency in 17 time periods, stratifying by sample location. We also retrieved the allele age estimate for rs1790218 using the Human Genome Dating tool^59^ with all default settings.

### Statistical Analysis

When comparing the significant differences among HEK293 cells transfected with GFP only and various SLC22A10 ortholog species or other transporters, we performed multiple comparisons using one-way analysis of variance followed by Dunnett’s two-tailed test. HEK293 cells transiently transfected with the GFP vector served as the control. The fold uptake of the substrate, relative to the control cells, was plotted based on one representative experiment conducted in triplicate wells (mean ± s.d.). Statistical significance was indicated as ***p<0.0001, **p<0.01, *p<0.05. These findings were further confirmed through at least one or two additional experiments. For specific differences and more detailed information, please refer to the figure legend.

## Funding/Acknowledgements

This research was supported by NIH GM117163 (SWY) and GM139875 (KMG, SWY). Additional support was provided by USDA Intramural Projects 2032-51530-025-00D (JWN). The USDA is an equal opportunity employer and provider. TMB is supported by funding from the European Research Council (ERC) under the European Union’s Horizon 2020 research and innovation program (grant agreement No. 864203), PID2021-126004NB-100 (MICIIN/FEDER, UE) and Secretaria d’Universitats i Recerca and CERCA Programme del Departament d’Economia i Coneixement de la Generalitat de Catalunya (GRC 2021 SGR 00177). We would like to acknowledge Esther Lizano, Alba Duch and Mina Azimi for their invaluable assistance and support in the successful execution of our research.

## References

1. Meixner E, Goldmann U, Sedlyarov V, Scorzoni S, Rebsamen M, Girardi E, Superti-Furga G. A substrate-based ontology for human solute carriers. Mol Syst Biol. 2020;16(7):e9652. Epub 2020/07/23. doi: 10.15252/msb.20209652. PubMed PMID: 32697042; PMCID: PMC7374931.

2. Yee SW, Buitrago D, Stecula A, Ngo HX, Chien HC, Zou L, Koleske ML, Giacomini KM. Deorphaning a solute carrier 22 family member, SLC22A15, through functional genomic studies. FASEB J. 2020;34(12):15734–52. Epub 2020/10/31. doi: 10.1096/fj.202001497R. PubMed PMID: 33124720; PMCID: PMC7839234.

3. Yee SW, Stecula A, Chien HC, Zou L, Feofanova EV, van Borselen M, Cheung KWK, Yousri NA, Suhre K, Kinchen JM, Boerwinkle E, Irannejad R, Yu B, Giacomini KM. Unraveling the functional role of the orphan solute carrier, SLC22A24 in the transport of steroid conjugates through metabolomic and genome-wide association studies. PLoS Genet. 2019;15(9):e1008208. Epub 2019/09/26. doi: 10.1371/journal.pgen.1008208. PubMed PMID: 31553721; PMCID: PMC6760779 competing interests exist.

4. Girardi E, Agrimi G, Goldmann U, Fiume G, Lindinger S, Sedlyarov V, Srndic I, Gurtl B, Agerer B, Kartnig F, Scarcia P, Di Noia MA, Lineiro E, Rebsamen M, Wiedmer T, Bergthaler A, Palmieri L, Superti-Furga G. Epistasis-driven identification of SLC25A51 as a regulator of human mitochondrial NAD import. Nat Commun. 2020;11(1):6145. Epub 2020/12/03. doi: 10.1038/s41467-020-19871-x. PubMed PMID: 33262325; PMCID: PMC7708531.

5. Higuchi K, Sugiyama K, Tomabechi R, Kishimoto H, Inoue K. Mammalian monocarboxylate transporter 7 (MCT7/Slc16a6) is a novel facilitative taurine transporter. J Biol Chem. 2022;298(4):101800. Epub 2022/03/09. doi: 10.1016/j.jbc.2022.101800. PubMed PMID: 35257743; PMCID: PMC8980330.

6. Yee SW, Giacomini KM. Emerging Roles of the Human Solute Carrier 22 Family. Drug Metab Dispos. 2021;50(9):1193–210. Epub 2021/12/19. doi: 10.1124/dmd.121.000702. PubMed PMID: 34921098; PMCID: PMC9488978.

7. Ren T, Jones RS, Morris ME. Untargeted metabolomics identifies the potential role of monocarboxylate transporter 6 (MCT6/SLC16A5) in lipid and amino acid metabolism pathways. Pharmacol Res Perspect. 2022;10(3):e00944. Epub 2022/04/26. doi: 10.1002/prp2.944. PubMed PMID: 35466588; PMCID: PMC9035569.

8. Lindberg FA, Nordenankar K, Forsberg EC, Fredriksson R. SLC38A10 Deficiency in Mice Affects Plasma Levels of Threonine and Histidine in Males but Not in Females: A Preliminary Characterization Study of SLC38A10(-/-) Mice. Genes (Basel). 2023;14(4). Epub 2023/04/28. doi: 10.3390/genes14040835. PubMed PMID: 37107593; PMCID: PMC10138244.

9. Giacomini KM, Yee SW, Koleske ML, Zou L, Matsson P, Chen EC, Kroetz DL, Miller MA, Gozalpour E, Chu X. New and Emerging Research on Solute Carrier and ATP Binding Cassette Transporters in Drug Discovery and Development: Outlook From the International Transporter Consortium. Clin Pharmacol Ther. 2022;112(3):540–61. Epub 2022/05/01. doi: 10.1002/cpt.2627. PubMed PMID: 35488474; PMCID: PMC9398938.

10. Sun W, Wu RR, van Poelje PD, Erion MD. Isolation of a family of organic anion transporters from human liver and kidney. Biochem Biophys Res Commun. 2001;283(2):417–22. Epub 2001/05/01. doi: 10.1006/bbrc.2001.4774. PubMed PMID: 11327718.

11. Uhlen M, Fagerberg L, Hallstrom BM, Lindskog C, Oksvold P, Mardinoglu A, Sivertsson A, Kampf C, Sjostedt E, Asplund A, Olsson I, Edlund K, Lundberg E, Navani S, Szigyarto CA, Odeberg J, Djureinovic D, Takanen JO, Hober S, Alm T, Edqvist PH, Berling H, Tegel H, Mulder J, Rockberg J, Nilsson P, Schwenk JM, Hamsten M, von Feilitzen K, Forsberg M, Persson L, Johansson F, Zwahlen M, von Heijne G, Nielsen J, Ponten F. Proteomics. Tissue-based map of the human proteome. Science. 2015;347(6220):1260419. Epub 2015/01/24. doi: 10.1126/science.1260419. PubMed PMID: 25613900.

12. Ruiz-Orera J, Hernandez-Rodriguez J, Chiva C, Sabido E, Kondova I, Bontrop R, Marques-Bonet T, Alba MM. Origins of De Novo Genes in Human and Chimpanzee. PLoS Genet. 2015;11(12):e1005721. Epub 2016/01/01. doi: 10.1371/journal.pgen.1005721. PubMed PMID: 26720152; PMCID: PMC4697840.

13. Wilks C, Zheng SC, Chen FY, Charles R, Solomon B, Ling JP, Imada EL, Zhang D, Joseph L, Leek JT, Jaffe AE, Nellore A, Collado-Torres L, Hansen KD, Langmead B. recount3: summaries and queries for large-scale RNA-seq expression and splicing. Genome Biol. 2021;22(1):323. Epub 2021/12/01. doi: 10.1186/s13059-021-02533-6. PubMed PMID: 34844637; PMCID: PMC8628444.

14. Tiemann JKS, Zschach H, Lindorff-Larsen K, Stein A. Interpreting the molecular mechanisms of disease variants in human transmembrane proteins. Biophys J. 2023;122(11):2176–91. Epub 2023/01/06. doi: 10.1016/j.bpj.2022.12.031. PubMed PMID: 36600598; PMCID: PMC10257119.

15. Cagiada M, Johansson KE, Valanciute A, Nielsen SV, Hartmann-Petersen R, Yang JJ, Fowler DM, Stein A, Lindorff-Larsen K. Understanding the Origins of Loss of Protein Function by Analyzing the Effects of Thousands of Variants on Activity and Abundance. Mol Biol Evol. 2021;38(8):3235–46. Epub 2021/03/30. doi: 10.1093/molbev/msab095. PubMed PMID: 33779753; PMCID: PMC8321532.

16. Zou L, Stecula A, Gupta A, Prasad B, Chien HC, Yee SW, Wang L, Unadkat JD, Stahl SH, Fenner KS, Giacomini KM. Molecular Mechanisms for Species Differences in Organic Anion Transporter 1, OAT1: Implications for Renal Drug Toxicity. Mol Pharmacol. 2018;94(1):689–99. Epub 2018/05/04. doi: 10.1124/mol.117.111153. PubMed PMID: 29720497; PMCID: PMC5987998.

17. Dvorak V, Wiedmer T, Ingles-Prieto A, Altermatt P, Batoulis H, Barenz F, Bender E, Digles D, Durrenberger F, Heitman LH, AP IJ, Kell DB, Kickinger S, Korzo D, Leippe P, Licher T, Manolova V, Rizzetto R, Sassone F, Scarabottolo L, Schlessinger A, Schneider V, Sijben HJ, Steck AL, Sundstrom H, Tremolada S, Wilhelm M, Wright Muelas M, Zindel D, Steppan CM, Superti-Furga G. An Overview of Cell-Based Assay Platforms for the Solute Carrier Family of Transporters. Front Pharmacol. 2021;12:722889. Epub 2021/08/28. doi: 10.3389/fphar.2021.722889. PubMed PMID: 34447313; PMCID: PMC8383457.

18. Kuang W, Zhang J, Lan Z, Deepak R, Liu C, Ma Z, Cheng L, Zhao X, Meng X, Wang W, Wang X, Xu L, Jiao Y, Luo Q, Meng Z, Kee K, Liu X, Deng H, Li W, Fan H, Chen L. SLC22A14 is a mitochondrial riboflavin transporter required for sperm oxidative phosphorylation and male fertility. Cell Rep. 2021;35(3):109025. Epub 2021/04/22. doi: 10.1016/j.celrep.2021.109025. PubMed PMID: 33882315; PMCID: PMC8065176.

19. Wegler C, Wisniewski JR, Robertsen I, Christensen H, Kristoffer Hertel J, Hjelmesaeth J, Jansson-Lofmark R, Asberg A, Andersson TB, Artursson P. Drug Disposition Protein Quantification in Matched Human Jejunum and Liver From Donors With Obesity. Clin Pharmacol Ther. 2022;111(5):1142–54. Epub 2022/02/15. doi: 10.1002/cpt.2558. PubMed PMID: 35158408; PMCID: PMC9310776.

20. Wegler C, Matsson P, Krogstad V, Urdzik J, Christensen H, Andersson TB, Artursson P. Influence of Proteome Profiles and Intracellular Drug Exposure on Differences in CYP Activity in Donor-Matched Human Liver Microsomes and Hepatocytes. Mol Pharm. 2021;18(4):1792–805. Epub 2021/03/20. doi: 10.1021/acs.molpharmaceut.1c00053. PubMed PMID: 33739838; PMCID: PMC8041379.

21. Niu L, Geyer PE, Gupta R, Santos A, Meier F, Doll S, Wewer Albrechtsen NJ, Klein S, Ortiz C, Uschner FE, Schierwagen R, Trebicka J, Mann M. Dynamic human liver proteome atlas reveals functional insights into disease pathways. Mol Syst Biol. 2022;18(5):e10947. Epub 2022/05/18. doi: 10.15252/msb.202210947. PubMed PMID: 35579278; PMCID: PMC9112488.

22. Eide Kvitne K, Hole K, Krogstad V, Wollmann BM, Wegler C, Johnson LK, Hertel JK, Artursson P, Karlsson C, Andersson S, Andersson TB, Sandbu R, Hjelmesaeth J, Skovlund E, Christensen H, Jansson-Lofmark R, Asberg A, Molden E, Robertsen I. Correlations between 4beta-hydroxycholesterol and hepatic and intestinal CYP3A4: protein expression, microsomal ex vivo activity, and in vivo activity in patients with a wide body weight range. Eur J Clin Pharmacol. 2022;78(8):1289–99. Epub 2022/06/02. doi: 10.1007/s00228-022-03336-9. PubMed PMID: 35648149; PMCID: PMC9283167.

23. El-Khateeb E, Al-Majdoub ZM, Rostami-Hodjegan A, Barber J, Achour B. Proteomic Quantification of Changes in Abundance of Drug-Metabolizing Enzymes and Drug Transporters in Human Liver Cirrhosis: Different Methods, Similar Outcomes. Drug Metab Dispos. 2021;49(8):610–8. Epub 2021/05/29. doi: 10.1124/dmd.121.000484. PubMed PMID: 34045218.

24. Cordes FS, Bright JN, Sansom MS. Proline-induced distortions of transmembrane helices. J Mol Biol. 2002;323(5):951–60. Epub 2002/11/06. doi: 10.1016/s0022-2836(02)01006-9. PubMed PMID: 12417206.

25. Karczewski KJ, Francioli LC, Tiao G, Cummings BB, Alfoldi J, Wang Q, Collins RL, Laricchia KM, Ganna A, Birnbaum DP, Gauthier LD, Brand H, Solomonson M, Watts NA, Rhodes D, Singer-Berk M, England EM, Seaby EG, Kosmicki JA, Walters RK, Tashman K, Farjoun Y, Banks E, Poterba T, Wang A, Seed C, Whiffin N, Chong JX, Samocha KE, Pierce-Hoffman E, Zappala Z, O’Donnell-Luria AH, Minikel EV, Weisburd B, Lek M, Ware JS, Vittal C, Armean IM, Bergelson L, Cibulskis K, Connolly KM, Covarrubias M, Donnelly S, Ferriera S, Gabriel S, Gentry J, Gupta N, Jeandet T, Kaplan D, Llanwarne C, Munshi R, Novod S, Petrillo N, Roazen D, Ruano-Rubio V, Saltzman A, Schleicher M, Soto J, Tibbetts K, Tolonen C, Wade G, Talkowski ME, Genome Aggregation Database C, Neale BM, Daly MJ, MacArthur DG. The mutational constraint spectrum quantified from variation in 141,456 humans. Nature. 2020;581(7809):434–43. Epub 2020/05/29. doi: 10.1038/s41586-020-2308-7. PubMed PMID: 32461654; PMCID: PMC7334197 from Takeda Pharmaceutical Company. A.H.O’D.-L. has received honoraria from ARUP and Chan Zuckerberg Initiative. E.V.M. has received research support in the form of charitable contributions from Charles River Laboratories and Ionis Pharmaceuticals, and has consulted for Deerfield Management. J.S.W. is a consultant for MyoKardia. B.M.N. is a member of the scientific advisory board at Deep Genomics and consultant for Camp4 Therapeutics, Takeda Pharmaceutical, and Biogen. M.J.D. is a founder of Maze Therapeutics. D.G.M. is a founder with equity in Goldfinch Bio, and has received research support from AbbVie, Astellas, Biogen, BioMarin, Eisai, Merck, Pfizer, and Sanofi-Genzyme. The views expressed in this article are those of the author(s) and not necessarily those of the NHS, the NIHR, or the Department of Health. M.I.M. has served on advisory panels for Pfizer, NovoNordisk, Zoe Global; has received honoraria from Merck, Pfizer, NovoNordisk and Eli Lilly; has stock options in Zoe Global and has received research funding from Abbvie, Astra Zeneca, Boehringer Ingelheim, Eli Lilly, Janssen, Merck, NovoNordisk, Pfizer, Roche, Sanofi Aventis, Servier & Takeda. As of June 2019, M.I.M. is an employee of Genentech, and holds stock in Roche. N.R. is a non-executive director of AstraZeneca.

26. Taliun D, Harris DN, Kessler MD, Carlson J, Szpiech ZA, Torres R, Taliun SAG, Corvelo A, Gogarten SM, Kang HM, Pitsillides AN, LeFaive J, Lee SB, Tian X, Browning BL, Das S, Emde AK, Clarke WE, Loesch DP, Shetty AC, Blackwell TW, Smith AV, Wong Q, Liu X, Conomos MP, Bobo DM, Aguet F, Albert C, Alonso A, Ardlie KG, Arking DE, Aslibekyan S, Auer PL, Barnard J, Barr RG, Barwick L, Becker LC, Beer RL, Benjamin EJ, Bielak LF, Blangero J, Boehnke M, Bowden DW, Brody JA, Burchard EG, Cade BE, Casella JF, Chalazan B, Chasman DI, Chen YI, Cho MH, Choi SH, Chung MK, Clish CB, Correa A, Curran JE, Custer B, Darbar D, Daya M, de Andrade M, DeMeo DL, Dutcher SK, Ellinor PT, Emery LS, Eng C, Fatkin D, Fingerlin T, Forer L, Fornage M, Franceschini N, Fuchsberger C, Fullerton SM, Germer S, Gladwin MT, Gottlieb DJ, Guo X, Hall ME, He J, Heard-Costa NL, Heckbert SR, Irvin MR, Johnsen JM, Johnson AD, Kaplan R, Kardia SLR, Kelly T, Kelly S, Kenny EE, Kiel DP, Klemmer R, Konkle BA, Kooperberg C, Kottgen A, Lange LA, Lasky-Su J, Levy D, Lin X, Lin KH, Liu C, Loos RJF, Garman L, Gerszten R, Lubitz SA, Lunetta KL, Mak ACY, Manichaikul A, Manning AK, Mathias RA, McManus DD, McGarvey ST, Meigs JB, Meyers DA, Mikulla JL, Minear MA, Mitchell BD, Mohanty S, Montasser ME, Montgomery C, Morrison AC, Murabito JM, Natale A, Natarajan P, Nelson SC, North KE, O’Connell JR, Palmer ND, Pankratz N, Peloso GM, Peyser PA, Pleiness J, Post WS, Psaty BM, Rao DC, Redline S, Reiner AP, Roden D, Rotter JI, Ruczinski I, Sarnowski C, Schoenherr S, Schwartz DA, Seo JS, Seshadri S, Sheehan VA, Sheu WH, Shoemaker MB, Smith NL, Smith JA, Sotoodehnia N, Stilp AM, Tang W, Taylor KD, Telen M, Thornton TA, Tracy RP, Van Den Berg DJ, Vasan RS, Viaud-Martinez KA, Vrieze S, Weeks DE, Weir BS, Weiss ST, Weng LC, Willer CJ, Zhang Y, Zhao X, Arnett DK, Ashley-Koch AE, Barnes KC, Boerwinkle E, Gabriel S, Gibbs R, Rice KM, Rich SS, Silverman EK, Qasba P, Gan W, Consortium NT-OfPM, Papanicolaou GJ, Nickerson DA, Browning SR, Zody MC, Zollner S, Wilson JG, Cupples LA, Laurie CC, Jaquish CE, Hernandez RD, O’Connor TD, Abecasis GR. Sequencing of 53,831 diverse genomes from the NHLBI TOPMed Program. Nature. 2021;590(7845):290–9. Epub 2021/02/12. doi: 10.1038/s41586-021-03205-y. PubMed PMID: 33568819; PMCID: PMC7875770.

27. Scott EM, Halees A, Itan Y, Spencer EG, He Y, Azab MA, Gabriel SB, Belkadi A, Boisson B, Abel L, Clark AG, Greater Middle East Variome C, Alkuraya FS, Casanova JL, Gleeson JG. Characterization of Greater Middle Eastern genetic variation for enhanced disease gene discovery. Nat Genet. 2016;48(9):1071–6. Epub 2016/07/19. doi: 10.1038/ng.3592. PubMed PMID: 27428751; PMCID: PMC5019950.

28. Cropp CD, Komori T, Shima JE, Urban TJ, Yee SW, More SS, Giacomini KM. Organic anion transporter 2 (SLC22A7) is a facilitative transporter of cGMP. Mol Pharmacol. 2008;73(4):1151–8. Epub 2008/01/25. doi: 10.1124/mol.107.043117. PubMed PMID: 18216183; PMCID: PMC2698938.

29. Zheng D, Gerstein MB. The ambiguous boundary between genes and pseudogenes: the dead rise up, or do they? Trends Genet. 2007;23(5):219–24. Epub 2007/03/27. doi: 10.1016/j.tig.2007.03.003. PubMed PMID: 17382428.

30. Mighell AJ, Smith NR, Robinson PA, Markham AF. Vertebrate pseudogenes. FEBS Lett. 2000;468(2-3):109–14. Epub 2000/02/29. doi: 10.1016/s0014-5793(00)01199-6. PubMed PMID: 10692568.

31. Zhang ZD, Frankish A, Hunt T, Harrow J, Gerstein M. Identification and analysis of unitary pseudogenes: historic and contemporary gene losses in humans and other primates. Genome Biol. 2010;11(3):R26. Epub 2010/03/10. doi: 10.1186/gb-2010-11-3-r26. PubMed PMID: 20210993; PMCID: PMC2864566.

32. Seal RL, Braschi B, Gray K, Jones TEM, Tweedie S, Haim-Vilmovsky L, Bruford EA. Genenames.org: the HGNC resources in 2023. Nucleic Acids Res. 2023;51(D1):D1003–D9. Epub 2022/10/17. doi: 10.1093/nar/gkac888. PubMed PMID: 36243972; PMCID: PMC9825485.

33. Cunningham F, Allen JE, Allen J, Alvarez-Jarreta J, Amode MR, Armean IM, Austine-Orimoloye O, Azov AG, Barnes I, Bennett R, Berry A, Bhai J, Bignell A, Billis K, Boddu S, Brooks L, Charkhchi M, Cummins C, Da Rin Fioretto L, Davidson C, Dodiya K, Donaldson S, El Houdaigui B, El Naboulsi T, Fatima R, Giron CG, Genez T, Martinez JG, Guijarro-Clarke C, Gymer A, Hardy M, Hollis Z, Hourlier T, Hunt T, Juettemann T, Kaikala V, Kay M, Lavidas I, Le T, Lemos D, Marugan JC, Mohanan S, Mushtaq A, Naven M, Ogeh DN, Parker A, Parton A, Perry M, Pilizota I, Prosovetskaia I, Sakthivel MP, Salam AIA, Schmitt BM, Schuilenburg H, Sheppard D, Perez-Silva JG, Stark W, Steed E, Sutinen K, Sukumaran R, Sumathipala D, Suner MM, Szpak M, Thormann A, Tricomi FF, Urbina-Gomez D, Veidenberg A, Walsh TA, Walts B, Willhoft N, Winterbottom A, Wass E, Chakiachvili M, Flint B, Frankish A, Giorgetti S, Haggerty L, Hunt SE, GR II, Loveland JE, Martin FJ, Moore B, Mudge JM, Muffato M, Perry E, Ruffier M, Tate J, Thybert D, Trevanion SJ, Dyer S, Harrison PW, Howe KL, Yates AD, Zerbino DR, Flicek P. Ensembl 2022. Nucleic Acids Res. 2022;50(D1):D988–D95. Epub 2021/11/19. doi: 10.1093/nar/gkab1049. PubMed PMID: 34791404; PMCID: PMC8728283.

34. Koleske ML, McInnes G, Brown JEH, Thomas N, Hutchinson K, Chin MY, Koehl A, Arkin MR, Schlessinger A, Gallagher RC, Song YS, Altman RB, Giacomini KM. Functional genomics of OCTN2 variants informs protein-specific variant effect predictor for Carnitine Transporter Deficiency. Proc Natl Acad Sci U S A. 2022;119(46):e2210247119. Epub 2022/11/08. doi: 10.1073/pnas.2210247119. PubMed PMID: 36343260; PMCID: PMC9674959.

35. Azimi M, Yee SW, Riselli A, Silva DB, Giacomini CP, Giacomini KM, Brett CM. Characterization of P-glycoprotein orthologs from human, sheep, pig, dog, and cat. J Vet Pharmacol Ther. 2023. Epub 2023/05/18. doi: 10.1111/jvp.13386. PubMed PMID: 37198956.

36. Wisniewski JR, Mann M. Consecutive proteolytic digestion in an enzyme reactor increases depth of proteomic and phosphoproteomic analysis. Anal Chem. 2012;84(6):2631–7. Epub 2012/02/14. doi: 10.1021/ac300006b. PubMed PMID: 22324799.

37. Rappsilber J, Ishihama Y, Mann M. Stop and go extraction tips for matrix-assisted laser desorption/ionization, nanoelectrospray, and LC/MS sample pretreatment in proteomics. Anal Chem. 2003;75(3):663–70. Epub 2003/02/15. doi: 10.1021/ac026117i. PubMed PMID: 12585499.

38. Wiśniewski JR. Label-Free Quantitative Analysis of Mitochondrial Proteomes Using the Multienzyme Digestion-Filter Aided Sample Preparation (MED-FASP) and “Total Protein Approach”. In: Mokranjac D, Perocchi F, editors. Mitochondria Methods in Molecular Biology. New York: Humana Press; 2017.

39. Wisniewski JR, Gaugaz FZ. Fast and sensitive total protein and Peptide assays for proteomic analysis. Anal Chem. 2015;87(8):4110–6. Epub 2015/04/04. doi: 10.1021/ac504689z. PubMed PMID: 25837572.

40. Tyanova S, Temu T, Cox J. The MaxQuant computational platform for mass spectrometry-based shotgun proteomics. Nat Protoc. 2016;11(12):2301–19. Epub 2016/11/04. doi: 10.1038/nprot.2016.136. PubMed PMID: 27809316.

41. Wisniewski JR, Rakus D. Multi-enzyme digestion FASP and the ‘Total Protein Approach’-based absolute quantification of the Escherichia coli proteome. J Proteomics. 2014;109:322–31. Epub 2014/07/27. doi: 10.1016/j.jprot.2014.07.012. PubMed PMID: 25063446.

42. Jumper J, Evans R, Pritzel A, Green T, Figurnov M, Ronneberger O, Tunyasuvunakool K, Bates R, Zidek A, Potapenko A, Bridgland A, Meyer C, Kohl SAA, Ballard AJ, Cowie A, Romera-Paredes B, Nikolov S, Jain R, Adler J, Back T, Petersen S, Reiman D, Clancy E, Zielinski M, Steinegger M, Pacholska M, Berghammer T, Bodenstein S, Silver D, Vinyals O, Senior AW, Kavukcuoglu K, Kohli P, Hassabis D. Highly accurate protein structure prediction with AlphaFold. Nature. 2021;596(7873):583-9. Epub 2021/07/16. doi: 10.1038/s41586-021-03819-2. PubMed PMID: 34265844; PMCID: PMC8371605 have filed non-provisional patent applications 16/701,070 and PCT/EP2020/084238, and provisional patent applications 63/107,362, 63/118,917, 63/118,918, 63/118,921 and 63/118,919, each in the name of DeepMind Technologies Limited, each pending, relating to machine learning for predicting protein structures. The other authors declare no competing interests.

43. Varadi M, Anyango S, Deshpande M, Nair S, Natassia C, Yordanova G, Yuan D, Stroe O, Wood G, Laydon A, Zidek A, Green T, Tunyasuvunakool K, Petersen S, Jumper J, Clancy E, Green R, Vora A, Lutfi M, Figurnov M, Cowie A, Hobbs N, Kohli P, Kleywegt G, Birney E, Hassabis D, Velankar S. AlphaFold Protein Structure Database: massively expanding the structural coverage of protein-sequence space with high-accuracy models. Nucleic Acids Res. 2022;50(D1):D439–D44. Epub 2021/11/19. doi: 10.1093/nar/gkab1061. PubMed PMID: 34791371; PMCID: PMC8728224.

44. Lomize AL, Todd SC, Pogozheva ID. Spatial arrangement of proteins in planar and curved membranes by PPM 3.0. Protein Sci. 2022;31(1):209–20. Epub 2021/10/31. doi: 10.1002/pro.4219. PubMed PMID: 34716622; PMCID: PMC8740824.

45. Frenz B, Lewis SM, King I, DiMaio F, Park H, Song Y. Prediction of Protein Mutational Free Energy: Benchmark and Sampling Improvements Increase Classification Accuracy. Front Bioeng Biotechnol. 2020;8:558247. Epub 2020/11/03. doi: 10.3389/fbioe.2020.558247. PubMed PMID: 33134287; PMCID: PMC7579412.

46. Laine E, Karami Y, Carbone A. GEMME: A Simple and Fast Global Epistatic Model Predicting Mutational Effects. Mol Biol Evol. 2019;36(11):2604–19. Epub 2019/08/14. doi: 10.1093/molbev/msz179. PubMed PMID: 31406981; PMCID: PMC6805226.

47. Ke Y, Gonthier R, Isabelle M, Bertin J, Simard JN, Dury AY, Labrie F. A rapid and sensitive UPLC-MS/MS method for the simultaneous quantification of serum androsterone glucuronide, etiocholanolone glucuronide, and androstan-3alpha, 17beta diol 17-glucuronide in postmenopausal women. J Steroid Biochem Mol Biol. 2015;149:146–52. Epub 2015/02/24. doi: 10.1016/j.jsbmb.2015.02.009. PubMed PMID: 25701608.

48. Kuderna LFK, Gao H, Janiak MC, Kuhlwilm M, Orkin JD, Bataillon T, Manu S, Valenzuela A, Bergman J, Rousselle M, Silva FE, Agueda L, Blanc J, Gut M, de Vries D, Goodhead I, Harris RA, Raveendran M, Jensen A, Chuma IS, Horvath JE, Hvilsom C, Juan D, Frandsen P, Schraiber JG, de Melo FR, Bertuol F, Byrne H, Sampaio I, Farias I, Valsecchi J, Messias M, da Silva MNF, Trivedi M, Rossi R, Hrbek T, Andriaholinirina N, Rabarivola CJ, Zaramody A, Jolly CJ, Phillips-Conroy J, Wilkerson G, Abee C, Simmons JH, Fernandez-Duque E, Kanthaswamy S, Shiferaw F, Wu D, Zhou L, Shao Y, Zhang G, Keyyu JD, Knauf S, Le MD, Lizano E, Merker S, Navarro A, Nadler T, Khor CC, Lee J, Tan P, Lim WK, Kitchener AC, Zinner D, Gut I, Melin AD, Guschanski K, Schierup MH, Beck RMD, Umapathy G, Roos C, Boubli JP, Rogers J, Farh KK, Marques Bonet T. A global catalog of whole-genome diversity from 233 primate species. Science. 2023;380(6648):906-13. Epub 2023/06/01 23:42. doi: 10.1126/science.abn7829. PubMed PMID: 37262161.

49. de Manuel M, Kuhlwilm M, Frandsen P, Sousa VC, Desai T, Prado-Martinez J, Hernandez-Rodriguez J, Dupanloup I, Lao O, Hallast P, Schmidt JM, Heredia-Genestar JM, Benazzo A, Barbujani G, Peter BM, Kuderna LF, Casals F, Angedakin S, Arandjelovic M, Boesch C, Kuhl H, Vigilant L, Langergraber K, Novembre J, Gut M, Gut I, Navarro A, Carlsen F, Andres AM, Siegismund HR, Scally A, Excoffier L, Tyler-Smith C, Castellano S, Xue Y, Hvilsom C, Marques-Bonet T. Chimpanzee genomic diversity reveals ancient admixture with bonobos. Science. 2016;354(6311):477-81. Epub 2016/10/30. doi: 10.1126/science.aag2602. PubMed PMID: 27789843; PMCID: PMC5546212.

50. Nater A, Mattle-Greminger MP, Nurcahyo A, Nowak MG, de Manuel M, Desai T, Groves C, Pybus M, Sonay TB, Roos C, Lameira AR, Wich SA, Askew J, Davila-Ross M, Fredriksson G, de Valles G, Casals F, Prado-Martinez J, Goossens B, Verschoor EJ, Warren KS, Singleton I, Marques DA, Pamungkas J, Perwitasari-Farajallah D, Rianti P, Tuuga A, Gut IG, Gut M, Orozco-terWengel P, van Schaik CP, Bertranpetit J, Anisimova M, Scally A, Marques-Bonet T, Meijaard E, Krutzen M. Morphometric, Behavioral, and Genomic Evidence for a New Orangutan Species. Curr Biol. 2017;27(22):3576–7. Epub 2017/11/22. doi: 10.1016/j.cub.2017.11.020. PubMed PMID: 29161551.

51. Prado-Martinez J, Sudmant PH, Kidd JM, Li H, Kelley JL, Lorente-Galdos B, Veeramah KR, Woerner AE, O’Connor TD, Santpere G, Cagan A, Theunert C, Casals F, Laayouni H, Munch K, Hobolth A, Halager AE, Malig M, Hernandez-Rodriguez J, Hernando-Herraez I, Prufer K, Pybus M, Johnstone L, Lachmann M, Alkan C, Twigg D, Petit N, Baker C, Hormozdiari F, Fernandez-Callejo M, Dabad M, Wilson ML, Stevison L, Camprubi C, Carvalho T, Ruiz-Herrera A, Vives L, Mele M, Abello T, Kondova I, Bontrop RE, Pusey A, Lankester F, Kiyang JA, Bergl RA, Lonsdorf E, Myers S, Ventura M, Gagneux P, Comas D, Siegismund H, Blanc J, Agueda-Calpena L, Gut M, Fulton L, Tishkoff SA, Mullikin JC, Wilson RK, Gut IG, Gonder MK, Ryder OA, Hahn BH, Navarro A, Akey JM, Bertranpetit J, Reich D, Mailund T, Schierup MH, Hvilsom C, Andres AM, Wall JD, Bustamante CD, Hammer MF, Eichler EE, Marques-Bonet T. Great ape genetic diversity and population history. Nature. 2013;499(7459):471-5. Epub 2013/07/05. doi: 10.1038/nature12228. PubMed PMID: 23823723; PMCID: PMC3822165.

52. Genomes Project C, Auton A, Brooks LD, Durbin RM, Garrison EP, Kang HM, Korbel JO, Marchini JL, McCarthy S, McVean GA, Abecasis GR. A global reference for human genetic variation. Nature. 2015;526(7571):68-74. Epub 2015/10/04. doi: 10.1038/nature15393. PubMed PMID: 26432245; PMCID: PMC4750478.

53. Mafessoni F, Grote S, de Filippo C, Slon V, Kolobova KA, Viola B, Markin SV, Chintalapati M, Peyregne S, Skov L, Skoglund P, Krivoshapkin AI, Derevianko AP, Meyer M, Kelso J, Peter B, Prufer K, Paabo S. A high-coverage Neandertal genome from Chagyrskaya Cave. Proc Natl Acad Sci U S A. 2020;117(26):15132–6. Epub 2020/06/18. doi: 10.1073/pnas.2004944117. PubMed PMID: 32546518; PMCID: PMC7334501.

54. Meyer M, Kircher M, Gansauge MT, Li H, Racimo F, Mallick S, Schraiber JG, Jay F, Prufer K, de Filippo C, Sudmant PH, Alkan C, Fu Q, Do R, Rohland N, Tandon A, Siebauer M, Green RE, Bryc K, Briggs AW, Stenzel U, Dabney J, Shendure J, Kitzman J, Hammer MF, Shunkov MV, Derevianko AP, Patterson N, Andres AM, Eichler EE, Slatkin M, Reich D, Kelso J, Paabo S. A high-coverage genome sequence from an archaic Denisovan individual. Science. 2012;338(6104):222-6. Epub 2012/09/01. doi: 10.1126/science.1224344. PubMed PMID: 22936568; PMCID: PMC3617501.

55. Prufer K, de Filippo C, Grote S, Mafessoni F, Korlevic P, Hajdinjak M, Vernot B, Skov L, Hsieh P, Peyregne S, Reher D, Hopfe C, Nagel S, Maricic T, Fu Q, Theunert C, Rogers R, Skoglund P, Chintalapati M, Dannemann M, Nelson BJ, Key FM, Rudan P, Kucan Z, Gusic I, Golovanova LV, Doronichev VB, Patterson N, Reich D, Eichler EE, Slatkin M, Schierup MH, Andres AM, Kelso J, Meyer M, Paabo S. A high-coverage Neandertal genome from Vindija Cave in Croatia. Science. 2017;358(6363):655-8. Epub 2017/10/07. doi: 10.1126/science.aao1887. PubMed PMID: 28982794; PMCID: PMC6185897.

56. Prufer K, Racimo F, Patterson N, Jay F, Sankararaman S, Sawyer S, Heinze A, Renaud G, Sudmant PH, de Filippo C, Li H, Mallick S, Dannemann M, Fu Q, Kircher M, Kuhlwilm M, Lachmann M, Meyer M, Ongyerth M, Siebauer M, Theunert C, Tandon A, Moorjani P, Pickrell J, Mullikin JC, Vohr SH, Green RE, Hellmann I, Johnson PL, Blanche H, Cann H, Kitzman JO, Shendure J, Eichler EE, Lein ES, Bakken TE, Golovanova LV, Doronichev VB, Shunkov MV, Derevianko AP, Viola B, Slatkin M, Reich D, Kelso J, Paabo S. The complete genome sequence of a Neanderthal from the Altai Mountains. Nature. 2014;505(7481):43-9. Epub 2013/12/20. doi: 10.1038/nature12886. PubMed PMID: 24352235; PMCID: PMC4031459.

57. Machiela MJ, Chanock SJ. LDlink: a web-based application for exploring population-specific haplotype structure and linking correlated alleles of possible functional variants. Bioinformatics. 2015;31(21):3555–7. Epub 2015/07/04. doi: 10.1093/bioinformatics/btv402. PubMed PMID: 26139635; PMCID: PMC4626747.

58. Mallick S, Micco A, Mah M, Ringbauer H, Lazaridis I, Olalde I, Patterson N, Reich D. The Allen Ancient DNA Resource (AADR): A curated compendium of ancient human genomes. bioRxiv. 2023. Epub 2023/04/18. doi: 10.1101/2023.04.06.535797. PubMed PMID: 37066305; PMCID: PMC10104067.

59. Albers PK, McVean G. Dating genomic variants and shared ancestry in population-scale sequencing data. PLoS Biol. 2020;18(1):e3000586. Epub 2020/01/18. doi: 10.1371/journal.pbio.3000586. PubMed PMID: 31951611; PMCID: PMC6992231 following competing interests: GM is a shareholder in and non-executive director of Genomics PLC, and is a partner in Peptide Groove LLP. PKA is a shareholder in and a director of BioMe Oxford Ltd.

