## Supplementary Tables for "Illuminating the Function of the Orphan Transporter, SLC22A10 in Humans and Other Primates"

**Table S1.** List of steroid and steroid glucuronide tested.

| Compound | Vendor | Catalog number |
|---|---|---|
| d3-estradiol-3$\alpha$-glucuronide | Toronto Research Chemical | E888017 |
| estradiol-3β-glucuronide | Sigma-Aldrich | E2127-5MG |
| estradiol-17β-glucuronide | Sigma-Aldrich | E1127-10mg |
| estrone-3β-glucuronide | Toronto Research Chemical | E889055 |
| testosterone-17β-glucuronide | Steraloids | A6980-000 |
| androstanediol-3$\alpha$-glucuronide | Steraloids | A1206-800 |
| etiocholanolone-3$\alpha$-glucuronide | Steraloids | A3625-000 |
| androsterone-3-glucuronide | Toronto Research Chemical | A637605 |
| testosterone | Sigma-Aldrich | T1500 |
| progesterone | Sigma-Aldrich | P0130 |

**Table S2**. Steroid and steroid glucuronide mass spectrometer acquisition parameters.

| compound | tR (min) | ESI Mode | Q1 (Da) | Q3 (Da) | DP | CE |
|---|---|---|---|---|---|---|
| d3-estradiol-3$\alpha$-glucuronide | 1.5 | (-) | 450.1 | 274.2 | -25 | -25 |
| estradiol-3β-glucuronide | 1.90 | (-) | 447.2 | 271.2 | -25 | -25 |
| estradiol-17β-glucuronide | 1.94 | (-) | 447.2 | 271.2 | -25 | -40 |
| estrone-3β-glucuronide | 2.00 | (-) | 445.2 | 269.2 | -25 | -20 |
| testosterone-17β-glucuronide | 2.58 | (+) | 465.3 | 271.2 | 130 | 10 |
| androstanediol-3$\alpha$-glucuronide | 3.44 | (-) | 467.1 | 291.2 | -100 | -35 |
| etiocholanolone-3$\alpha$-glucuronide | 3.45 | (-) | 465.3 | 113.1 | -25 | -37 |
| androsterone-3-glucuronide | 3.46 | (-) | 465.3 | 85.2 | -25 | -40 |
| testosterone | 3.76 | (+) | 289.3 | 109.1 | 50 | 25 |
| 1-cycloheyl-3-uriedo-decanoic acid | 4.00 | (+) | 341.2 | 216.2 | 135 | 28 |
| 1-cycloheyl-3-uriedo-decanoic acid | 4.00 | (-) | 339.2 | 214.2 | -140 | -38 |
| progesterone | 4.20 | (+) | 315.2 | 109.1 | 40 | 28 |

The API 4500 source was operated at 650ºC, with spray voltage of 4500 V, curtain gas at 34 L/min, Gas 1at 50 L/min, Gas 2 at 20 L/min, and a collision gas setting of medium using nitrogen.
