## Supplementary Figures for "Illuminating the Function of the Orphan Transporter, SLC22A10 in Humans and Other Primates"

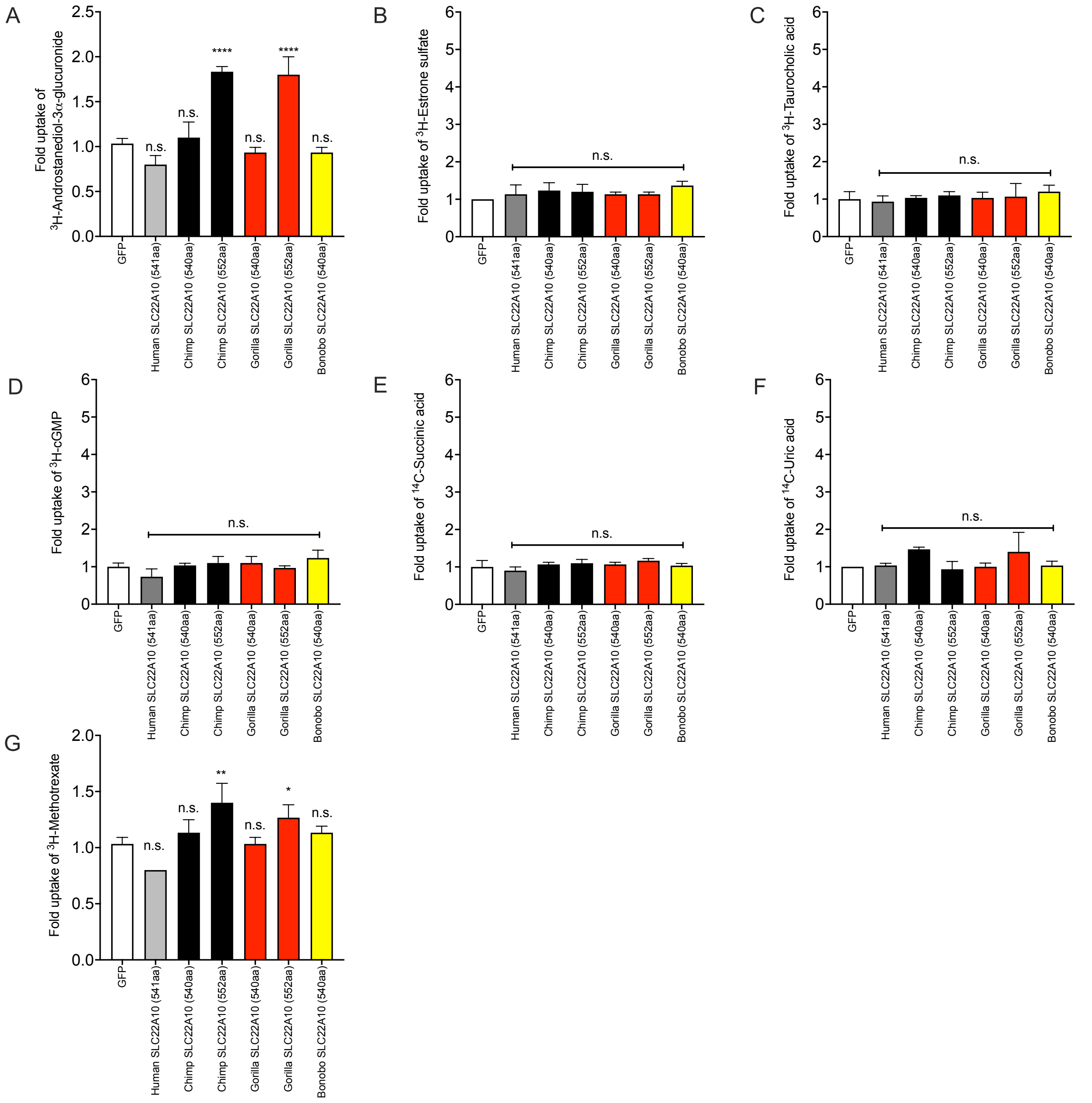


**Supplemental Figure 1**. Six anions were screened as substrates of SLC22A10. (A) [$^{3}$H]-androstanediol-3$\alpha$-glucuronide; (B) [$^{3}$H]-estrone sulfate; (C) [$^{3}$H]-taurocholic acid; (D) [$^{3}$H]-cGMP; (E) [$^{14}$C]-succinic acid, (F) [$^{14}$C]-uric acid, (G) [$^{3}$H]-methotrexate. Multiple comparisons were analyzed using one-way analysis of variance followed by Dunnett's two-tailed test. HEK293 cells transiently transfected with GFP vector was used as control. All expression vectors used have GFP-tagged in the N-terminal. One-Way Multiple comparisons were used to compare the mean of each orthologs with the mean of the control (GFP). Data are from one representative experiment in triplicate wells (mean ± s.d.). ****$p<0.0001$, ***$p<0.0005$, **$p<0.01$, *$p<0.05$. Results were replicated in at least one additional experiment.

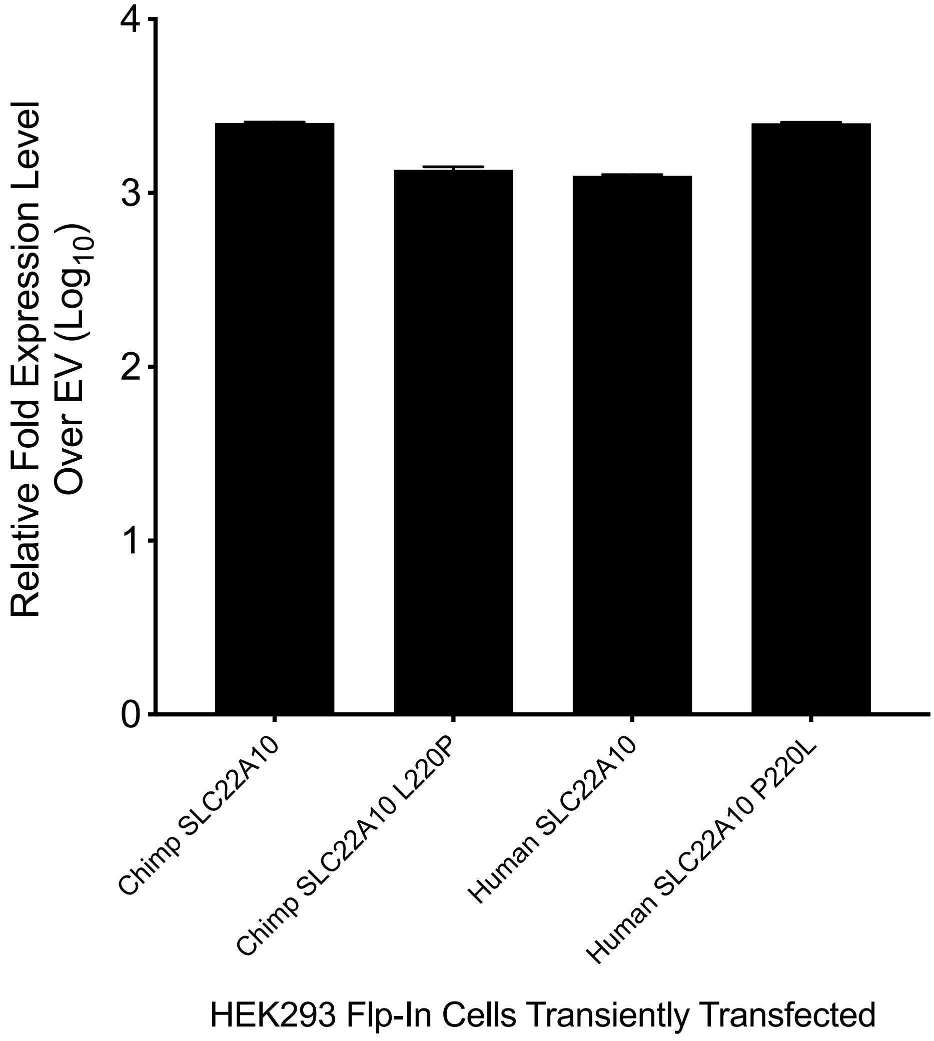


**Supplemental Figure 2**. Transcript levels of SLC22A10 in HEK293 cells that were transiently transfected with human and chimpanzee SLC22A10, as well as their respective mutations.

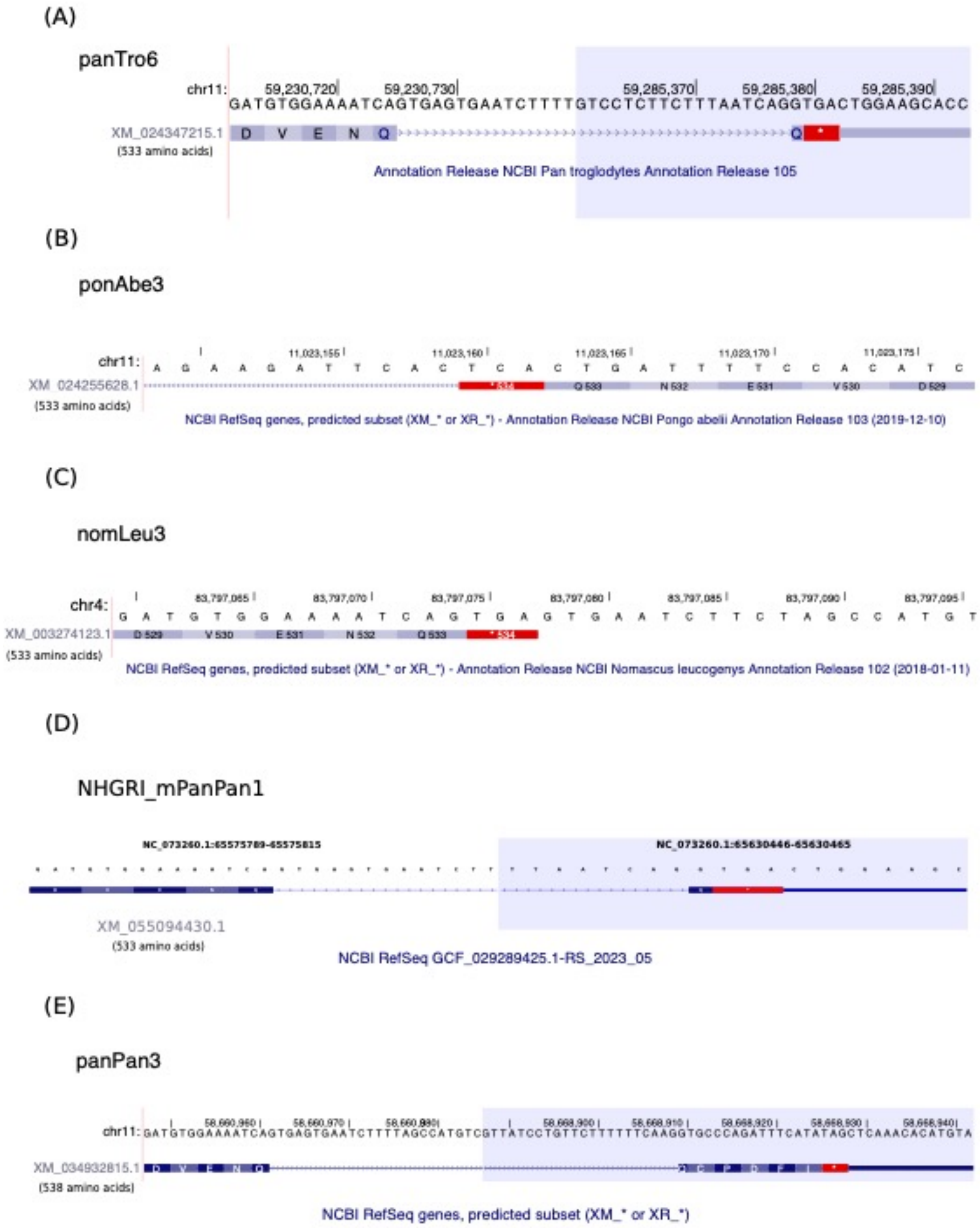


**Supplementary Figure 3.** SLC22A10 isoforms encoding 533 amino acids in chimpanzee, orangutan, gibbon and bonobo. In addition to the 552 amino acids SLC22A10, NCBI annotations show that chimpanzee (A), orangutan (B), and gibbon (C) and bonobo (D) are predicted to have isoforms with 533 amino acids. It is worth noting that at the beginning of this project, NCBI had initially predicted the bonobo SLC22A10 isoform to have 538 amino acids (panPan3 assembly) (E). These shorter protein isoforms are derived from alternative acceptor sites (chimpanzee/bonobo) and exon extensions (orangutan/gibbon). Blue shadows indicate a split view of the intron sequence between two exons.

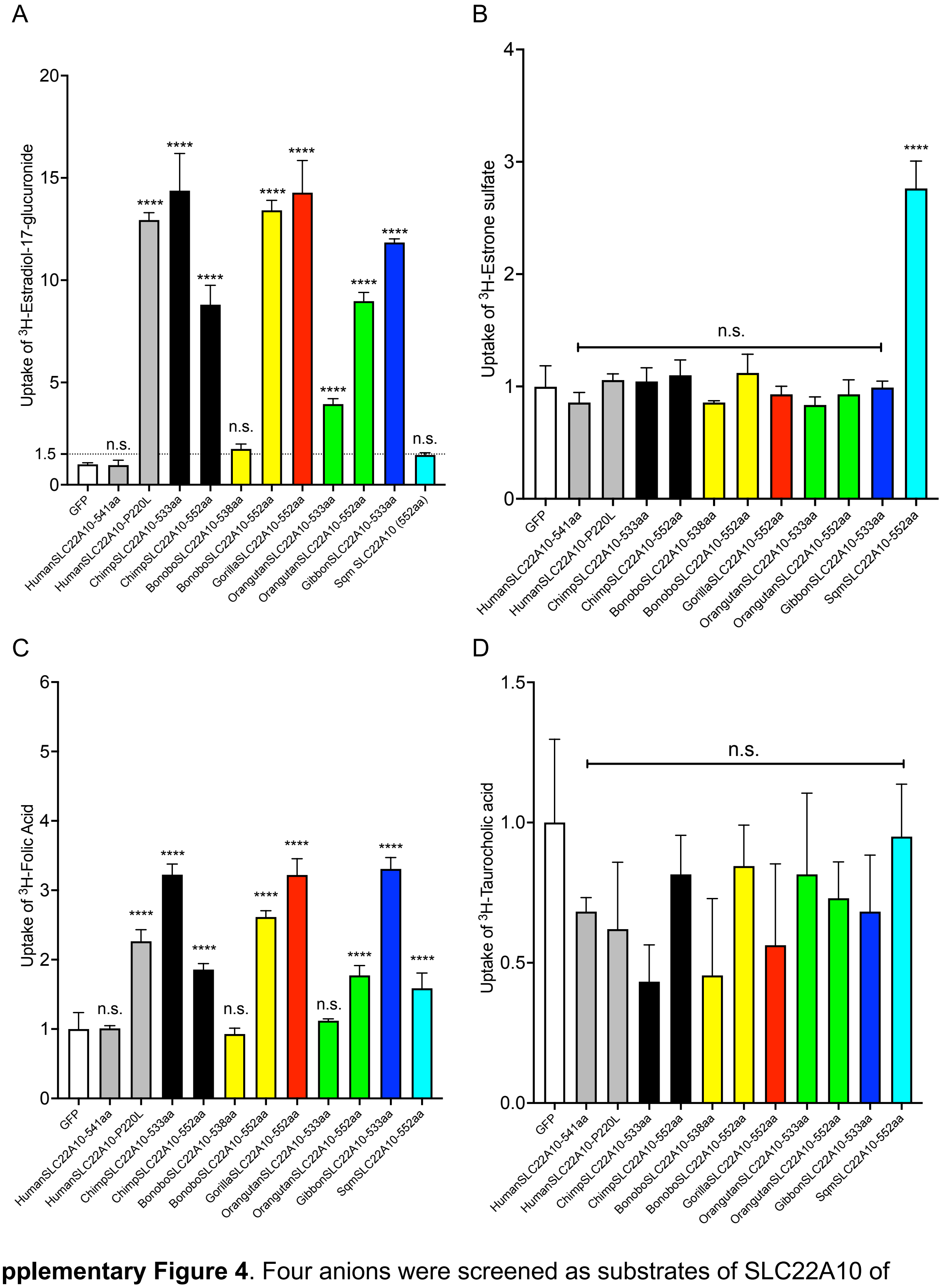


**Supplementary Figure 4**. Four anions were screened as substrates of SLC22A10 of different species and isoforms. Human SLC22A10 and human SLC22A10-P220L are also included in the assay. A. [$^3$H]-estradiol-17β-glucuronide; B. [$^3$H]-estrone sulfate; C. [$^3$H]-folic acid; and D. [$^3$H]-taurocholic acid. Multiple comparisons were analyzed using one-way analysis of variance followed by Dunnett's two-tailed test. HEK293 cells transiently transfected with GFP vector was used as control. Data are from one representative experiment in triplicate wells (mean ± s.d.). All expression vectors do not have GFP-tagged. Results was replicated in two independent experiments. n.s. Not significance compared to HEK293 cells transfected with GFP only. One-Way Multiple comparisons were used to compare the mean of each orthologs with the mean of the control (GFP). ****$p<0.0001$, ***$p<0.0005$, **$p<0.01$, *$p<0.05$. Results were replicated in at least one additional experiment.

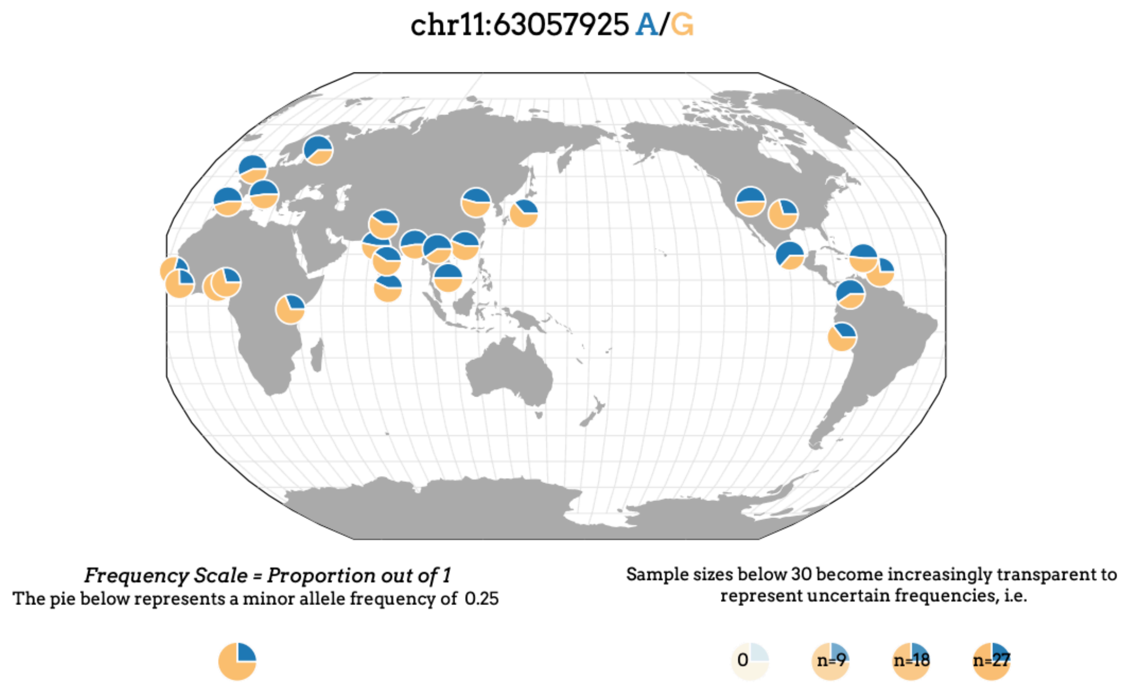


https://popgen.uchicago.edu/ggv/?data=%221000genomes%22&chr=11&pos=63057925

## 11-63290453-G-A

| | |
|---|---|
| Chromosome | 11 |
| Position | 63,290,453 |
| Reference allele | G |
| Alternate allele | A |
| rsID | rs1790218 |
| Filter | PASS |
| ClinVar | None |
| PubMed | None |

| | |
|---|---|
| Samples | 132,345 |
| AC (Alternate allele Count) | 123,421 |
| AF (Alternate allele Frequency) | 0.46628 |
| Heterozygotes | 61,911 |
| Homozygotes | 30,755 |

**Allele frequency in 1000G**

| | |
|---|---|
| AFR (African) | 0.2663 |
| ALL (All individuals) | 0.4335 |
| AMR (Ad Mixed American) | 0.5058 |
| EAS (East Asian) | 0.4692 |
| EUR (European) | 0.5537 |
| SAS (South Asian) | 0.4479 |

**Allele frequency in gnomAD r2.1**

| | |
|---|---|
| AFR (African) | 0.2887 |
| ALL (All individuals) | 0.5175 |
| AMR (Ad Mixed American) | 0.5078 |
| ASJ (Ashkenazi Jewish) | 0.4441 |
| EAS (East Asian) | 0.4216 |
| FIN (Finnish) | 0.6062 |
| NFE (Non-Finnish European) | 0.5754 |
| OTH (Others) | 0.5296 |
| SAS (South Asian) | 0.4521 |

**Supplementary Figure 5.** The distribution of the allele frequency of rs1790218 (SLC22A10-Trp96STOP) varies across different human populations. The rs1790218 variant (G>A) is a nonsense variant that encodes the A-allele and leads to a premature stop codon, resulting in the p.Trp96Ter alteration. The frequency of the A-allele varies, with approximately 28% in African populations and up to 60% in Finnish populations.

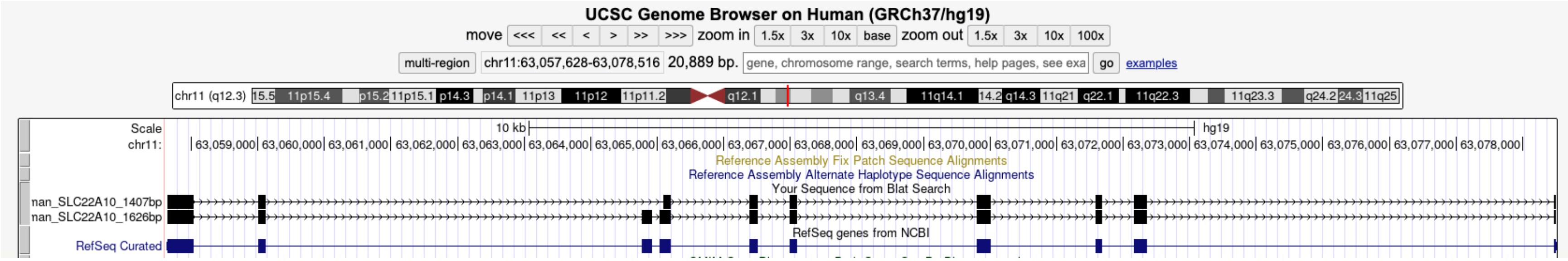


>human SLC22A10 open reading frame (three out of 27 colonies have the full SLC22A10 ORF)
ATGGCCTTTGAGGAGCTCTTGAGTCAAGTTGGAGGCCTTGGGAGATTTCAGATGCTTCATCTGGTTTTTATTCTTCCCTCTCTCAT
GTTATTAATCCCTCATATACTGCTAGAGAACTTTGCTGCAGCCATTCCTGGTCATCGTTGCTGGGTCCACATGCTGGACAATAATA
CTGGATCTGGTAATGAAACTGGAATCCTCAGTGAAGATGCCCTCTTGAGAATCTCTATCCCACTAGACTCAAATCTGAGGCCAGA
GAAGTGTCGTCGCTTTGTCCATCCCCAGTGGCAGCTTCTTCACCTGAATGGGACTATCCACAGCACAAGTGAGGCAGACACAGA
ACCCTGTGTGGATGGCTGGGTATATGATCAAAGCTACTTCCCTTCGACCATTGTGACTAAGTGGGACCTGGTATGTGATTATCAG
TCACTGAAATCAGTGGTTCAATTCCTACTTCTGACTGGAATGCTGGTGGGAGGCATCATAGGTGGCCATGTCTCAGACAGGTTTG
GGCGAAGATTTATTCTCAGATGGTGTTTGCTCCAGCTTGCCATTACTGACACCTGCGCTGCCTTCGCTCCCACCTTCCCTGTTTA
CTGTGTACTACGCTTCTTGGCAGGTTTTTCTTCCATGATCATTATATCAAATAATTCTTTGCCCATTACTGAGTGGATAAGGCCCA
ACTCTAAAGCCCTGGTAGTAATATTGTCATCTGGTGCCCTTAGTATTGGACAGATAATCCTGGGAGGCTTGGCTTATGTCTTCCG
AGACTGGCAAACCCTGCACGTGGTGGCGTCTGTACCTTTCTTTGTCTTCTTTCTTCTTTCAAGGTGGCTGGTGGAATCTGCTCGG
TGGTTGATAATCACCAATAAACTAGATGAGGGCTTAAAGGCACTTAGAAAAGTTGCACGCACAAATGGAATAAAGAATGCTGAAG
AAACCCTGAACATAGAGGTTGTAAGATCCACCATGCAGGAGGAGCTGGATGCAGCACAGACCAAAACTACTGTGTGTGACTTGT
TCCGCAACCCCAGTATGCGTAAAAGGATCTGTATCCTGGTATTTTTGAGATTTGCAAACACAATACCTTTTTATGGTACCATGGTC
AATCTTCAGCATGTGGGGAGCAACATTTTCCTGTTGCAGGTACTTTATGGAGCTGTCGCTCTCATAGTTCGATGTCTTGCTCTTTT
GACACTAAATCATATGGGCCGTCGAATAAGCCAGATATTGTTCATGTTCCTGGTGGGCCTTTCCATTTTGGCCAACACGTTTGTG
CCCAAAGAAATGCAGACCCTGCGTGTGGCTTTGGCATGTCTGGGAATCGGCTGTTCTGCTGCTACTTTTTCCAGTGTTGCTGTTC
ACTTCATTGAACTCATCCCCACTGTTCTCAGGGCAAGAGCTTCAGGAATAGATTTAACGGCTAGTAGGATTGGAGCAGCACTGG
CTCCCCTCTTGATGACCTTAACGGTATTTTTTACCACTTTGCCATGGATCATTTATGGAATCTTCCCCATCATTGGTGGCCTTATT
GTCTTCCTCCTACCAGAAACCAAGAATCTGCCTTTGCCTGACACCATCAAGGATGTGGAAAATCAAAAAAAAAATCTCAAGGAAA
AGGCATAA

>human SLC22A10 open reading frame (23 out of 27 colonies have the 219-bp deletion (in red) of the SLC22A10 ORF)
ATGGCCTTTGAGGAGCTCTTGAGTCAAGTTGGAGGCCTTGGGAGATTTCAGATGCTTCATCTGGTTTTTATTCTTCCCTCTCTCAT
GTTATTAATCCCTCATATACTGCTAGAGAACTTTGCTGCAGCCATTCCTGGTCATCGTTGCTGGGTCCACATGCTGGACAATAATA
CTGGATCTGGTAATGAAACTGGAATCCTCAGTGAAGATGCCCTCTTGAGAATCTCTATCCCACTAGACTCAAATCTGAGGCCAGA
GAAGTGTCGTCGCTTTGTCCATCCCCAGTGGCAGCTTCTTCACCTGAATGGGACTATCCACAGCACAAGTGAGGCAGACACAGA

ACCCTGTGTGGATGGCTGGGTATATGATCAAAGCTACTTCCCTTCGACCATTGTGACTAAGTGGGACCTGGTATGTGATTATCAG
TCACTGAAATCAGTGGTTCAATTCCTACTTCTGACTGGAATGCTGGTGGGAGGCATCATAGGTGGCCATGTCTCAGACAGGTTTG
GGCGAAGATTTATTCTCAGATGGTGTTTGCTCCAGCTTGCCATTACTGACACCTGCGCTGCCTTCGCTCCCACCTTCCCTGTTTA
CTGTGTACTACGCTTCTTGGCAGGTTTTTCTTCCATGATCATTATATCAAATAATTCTTTGCCCATTACTGAGTGGATAAGGCCCA
ACTCTAAAGCCCTGGTAGTAATATTGTCATCTGGTGCCCTTAGTATTGGACAGATAATCCTGGGAGGCTTGGCTTATGTCTTCCG
AGACTGGCAAACCCTGCACGTGGTGGCGTCTGTACCTTTCTTTGTCTTCTTTCTTCTTTCAAGGTGGCTGGTGGAATCTGCTCGG
TGGTTGATAATCACCAATAAACTAGATGAGGGCTTAAAGGCACTTAGAAAAGTTGCACGCACAAATGGAATAAAGAATGCTGAAG
AAACCCTGAACATAGAGGTTGTAAGATCCACCATGCAGGAGGAGCTGGATGCAGCACAGACCAAAACTACTGTGTGTGACTTGT
TCCGCAACCCCAGTATGCGTAAAAGGATCTGTATCCTGGTATTTTTGAGATTTGCAAACACAATACCTTTTTATGGTACCATGGTC
AATCTTCAGCATGTGGGGAGCAACATTTTCCTGTTGCAGGTACTTTATGGAGCTGTCGCTCTCATAGTTCGATGTCTTGCTCTTTT
GACACTAAATCATATGGGCCGTCGAATAAGCCAGATATTGTTCATGTTCCTGGTGGGCCTTTCCATTTTGGCCAACACGTTTGTG
CCCAAAGAAATGCAGACCCTGCGTGTGGCTTTGGCATGTCTGGGAATCGGCTGTTCTGCTGCTACTTTTTCCAGTGTTGCTGTTC
ACTTCATTGAACTCATCCCCACTGTTCTCAGGGCAAGAGCTTCAGGAATAGATTTAACGGCTAGTAGGATTGGAGCAGCACTGG
CTCCCCTCTTGATGACCTTAACGGTATTTTTTACCACTTTGCCATGGATCATTTATGGAATCTTCCCCATCATTGGTGGCCTTATT
GTCTTCCTCCTACCAGAAACCAAGAATCTGCCTTTGCCTGACACCATCAAGGATGTGGAAAATCAAAAAAAAAATCTCAAGGAAA
AGGCATAA

**Supplemental Figure 6.** Human SLC22A10 open reading frame (ORF). The first fasta sequence above is human SLC22A10 ORF with 1626 bp (NM_001039752). The second fasta sequence has a 219-bp deletion (in red), result in 1407 bp. The two fasta sequence were pasted in UCSC genome browser (hg19) to Blat search. The human SLC22A10 ORF with 1407bp skip the entire exon 3 and early part of exon 4.

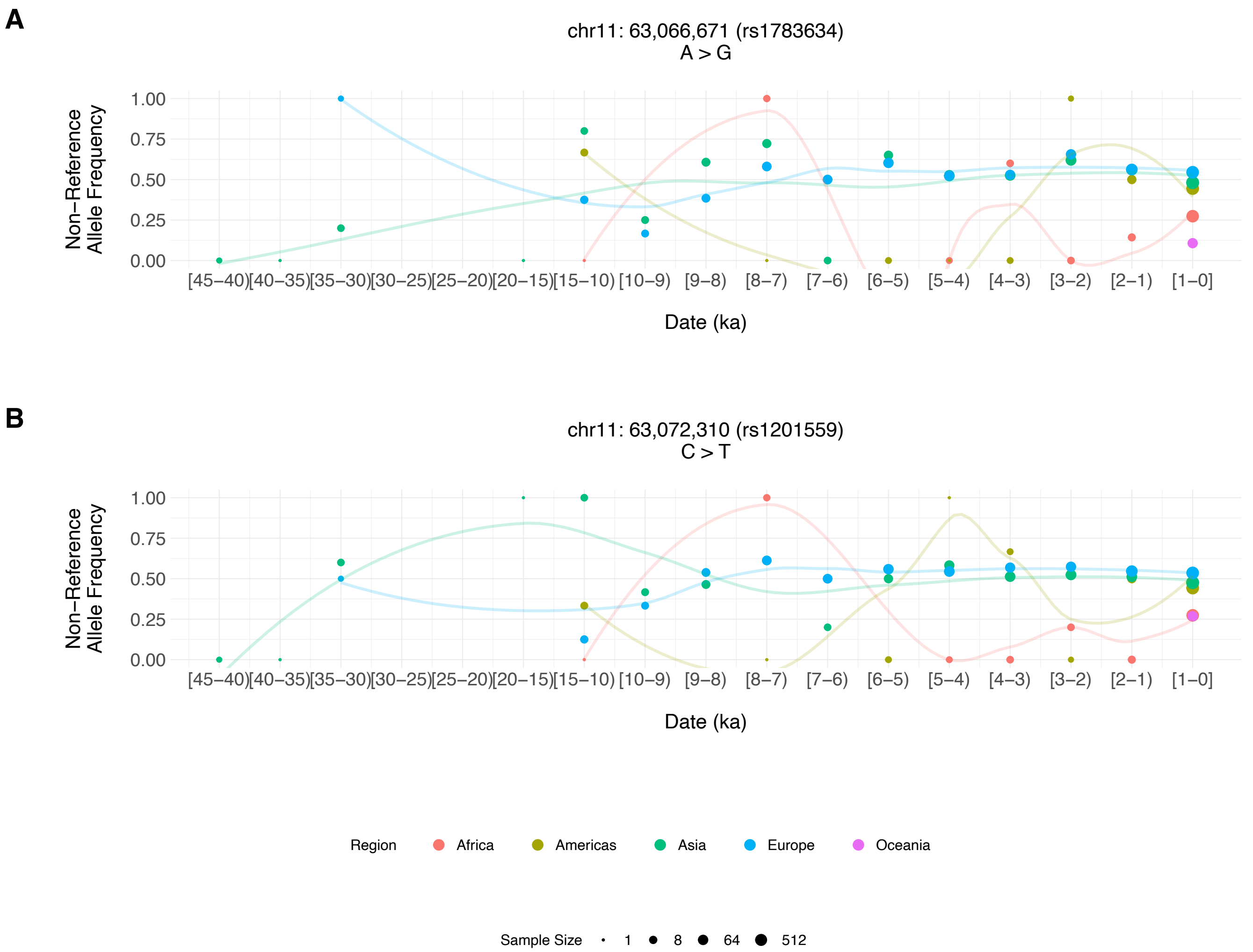


**Supplementary Figure 7**. Allele frequencies of two single nucleotide polymorphisms (SNPs), rs1783634 and rs1201559, were analyzed in diverse human populations. These SNPs are highly correlated (D' > 0.9, r2 > 0.9) with SLC22A10-Trp96Ter (rs1790218). The allele frequencies for these two SNPs were obtained from The Simons Genome Diversity Project (https://www.simonsfoundation.org/simons-genome-diversity-project/ ), 1000 Genomes Project, and the Allen Ancient DNA Resources (AADR) (https://reich.hms.harvard.edu/allen-ancient-dna-resource-aadr-downloadable-genotypes-present-day-and-ancient-dna-data ). The figure displays the allele frequencies of the alleles strongly linked to SLC22A10-96Ter.

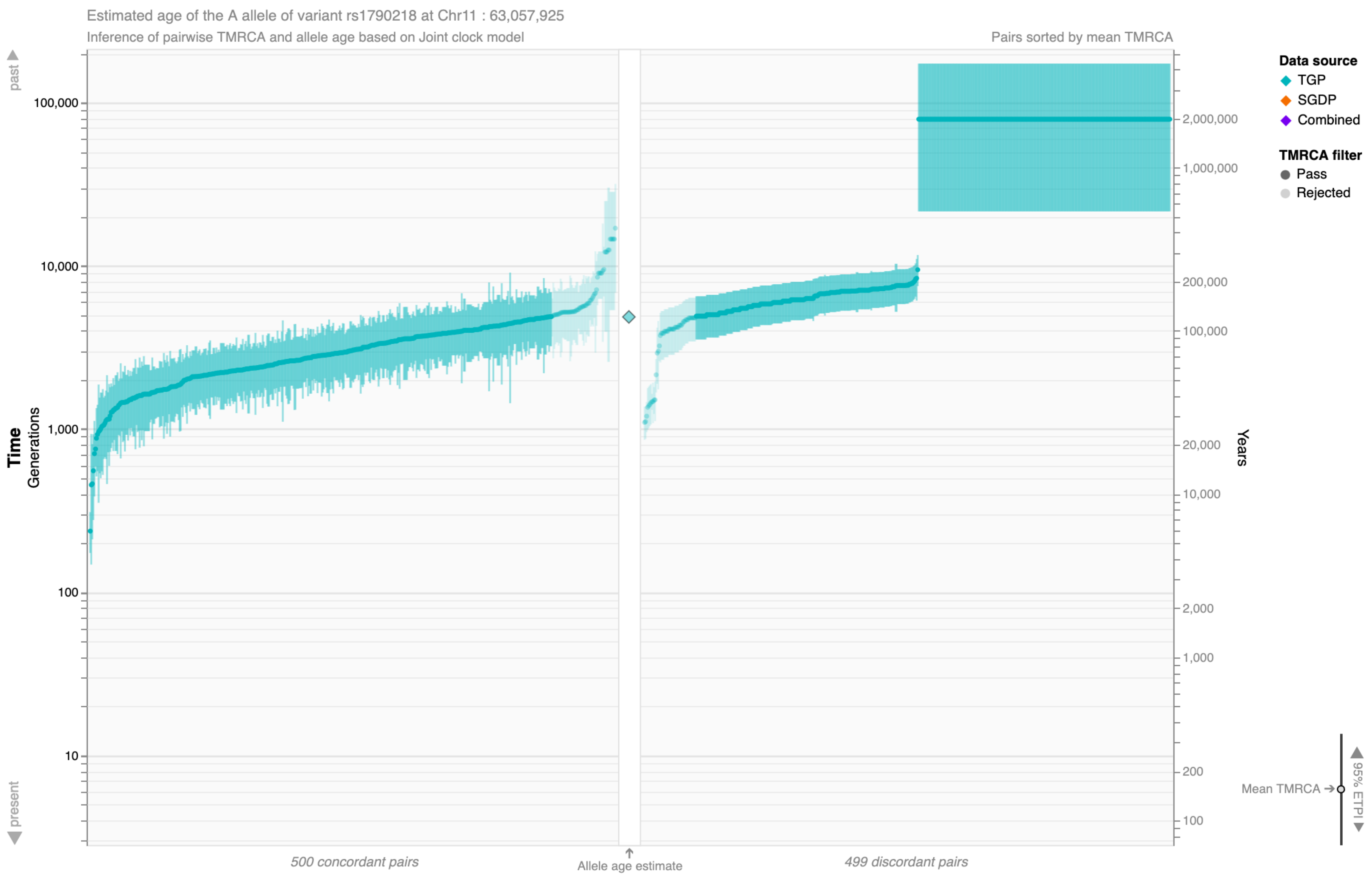
Estimated age of the A allele of variant rs1790218 at Chr11 : 63,057,925
Inference of pairwise TMRCA and allele age based on Joint clock model
Pairs sorted by mean TMRCA
past
present
Time
Generations
Years
100,000
10,000
1,000
100
10
2,000,000
1,000,000
200,000
100,000
20,000
10,000
2,000
1,000
200
100
500 concordant pairs
Allele age estimate
499 discordant pairs
Data source
TGP
SGDP
Combined
TMRCA filter
Pass
Rejected
Mean TMRCA
95% ETPI

**Supplementary Figure 8**. Allele age estimate for rs1790218 (G > A) from the Human Genome Dating portal using the joint clock model. The A allele is estimated to emerge 4,873 generations or 121,825 years ago (quality score = 0.878). ETPI = equal-tailed probability interval (95% credible interval), SGDP = Simons Genome Diversity Project, TGP = 1000 Genomes Project, TMRCA = time to most recent common ancestor.
